# CytoPheno: Automated descriptive cell type naming in flow and mass cytometry

**DOI:** 10.1101/2025.03.11.639902

**Authors:** Amanda R. Tursi, Celine S. Lages, Kenneth Quayle, Zachary T. Koenig, Rashi Loni, Shruti Eswar, José Cobeña-Reyes, Sherry Thornton, Tamara Tilburgs, Sandra Andorf

## Abstract

Advances in cytometry have led to increases in the number of cellular markers that are routinely measured. The resulting complexity of the data has prompted a shift from manual to automated analysis methods. Currently, numerous unsupervised methods are available to cluster cells based on marker expression values. However, phenotyping the resulting clusters is typically not part of the automated process. Manually identifying both marker definitions (e.g. CD4^+^, CCR7^+^, CD45RA^+^, CD19^-^) and descriptive cell type names (e.g. naïve CD4^+^ T cells) based on marker expression values can be time-consuming, subjective, and error-prone.

In this work we propose an algorithm that addresses these problems through the creation of an automated tool, CytoPheno, that assigns marker definitions and cell type names to unidentified clusters. First, post-clustered expression data undergoes per-marker calculations to assign markers as positive or negative. Next, marker names undergo a standardization process to match to Protein Ontology identifier terms. Finally, marker descriptions are matched to cell type names within the Cell Ontology. Each part of the tool was tested with benchmark data to demonstrate performance. Additionally, the tool is encompassed in a graphical user interface (R Shiny) to increase user accessibility and interpretability. Overall, CytoPheno can aid researchers in timely and unbiased phenotyping of post-clustered cytometry data.

## Introduction

Flow cytometry is used to characterize single cells in heterogeneous populations by staining cells with fluorescently labeled antibodies^1^. Similarly, mass cytometry is used to characterize cells with the usage of heavy metal isotope tagged antibodies^2^. Traditionally, the identification of cell populations is achieved by manual gating on a series of biaxial plots. However, manual gating has proven to be subjective, with studies showing inter-laboratory variability in gating strategies and the resulting cell types^3–7^. Furthermore, manual gating can be prohibitively time-consuming since the number of biaxial plots grows exponentially with dimensionality^8^. Advancements in both fluorescence and mass-based flow cytometry now allow for panels with over 40 markers, making manual analysis increasingly unfeasible^9–13^. Indeed, laboratories have reported that a single moderately-sized cytometry study requires 10 to 20 hours of analysis^14,15^.

With manual gating no longer as feasible, a shift towards automated analysis methods has occurred. Multiple unsupervised approaches have been developed to cluster cells based on their marker expression values^16–20^. Compared to manual gating, unsupervised clustering is less actively time-consuming and not limited by preconceived gating strategies^14,21^. As a result, it is good for novel population discovery and offers reproducibility advantages^14,21,22^. However, post-clustering phenotyping can still be laborious, as it largely depends on manual comparisons of median or mean marker expression levels and various visualization techniques (e.g., heatmaps, density plots, dimensionality reduction plots). For example, one study noted that for a 40 dimensional panel, phenotyping a single cell cluster amongst 10 total requires up to 360 comparisons^23^. Additionally, relying only on this type of manual approach has proved to be faulty and biased^23,24^. Even after the process of describing each marker per cell type is complete, the final step of assigning descriptive cell type names must be done. This can be an even more error-prone and subjective task since it is largely dependent on an analyst’s immunology knowledge and can unconsciously be swayed by expected results. Consulting a marker-cell type reference may limit these challenges, but the time-consuming problem remains.

This phenotyping bottleneck is better addressed by the use of semi-supervised or supervised methods^25–31^. These algorithms incorporate cell population information to cluster and annotate cells. However, despite this clear advantage, supervised methods have several downsides. They typically require more user input, are labor intensive, and can be difficult to implement^21^. Furthermore, they are limited in their ability to identify novel or rare cell types^21,32^. Consequently, unsupervised methods are still widely used for automated data analysis^33^.

This cytometry phenotyping annotation tool (CytoPheno) was created so scientists can retain the advantages of unsupervised methods while its major shortcoming – a lack of cellular automated phenotyping – is addressed (Figure 1). The tool fully encompasses phenotype identification of unknown cell clusters, allowing for the marker descriptions (Part 1), marker names (Part 2), and cell type names (Part 3) to be determined through a standardized multi-step process. Part 1 relies on categorizing individual markers used in the experiment as positive (expressed) or negative (not expressed) for each cluster. Part 2 standardizes marker names using various resources to ultimately output matched Protein Ontology (PRO) or Gene Ontology (GO) terms^34–36^. Part 3 translates the marker descriptions determined in Part 1 and the standardized marker names determined in Part 2 into descriptive cell type names from the Cell Ontology (CL) or Provisional Cell Ontology (PCL) that can be further interpreted by an immunologist^37^.

**Figure 1.**
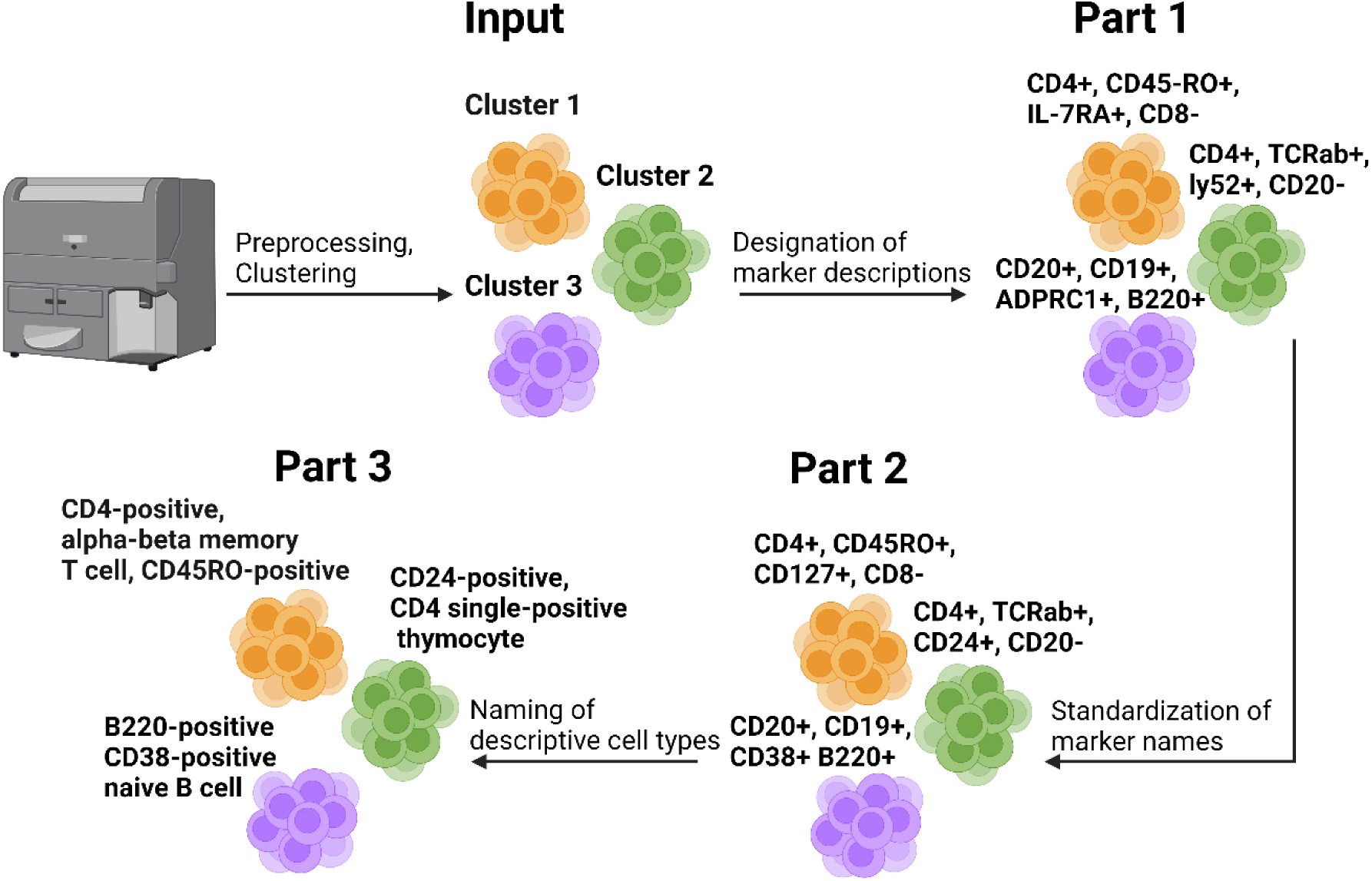
Simplified schematic depicting the inputs and outputs for each part of the CytoPheno tool. Created in BioRender.com.

Currently only a few methods for assigning marker definitions after unsupervised clustering exist, all of which are standalone tools that were designed with different purposes than that of this tool^23,38–40^. These tools aim to find the ‘most important’ markers per cell type, such as markers that are highly variable across cell subsets or markers that are optimal in assigning a sparse gating strategy. These algorithms may achieve their stated goals but are not ideal for the purpose of ultimately assigning cell type names. Existing methods for the automated assignment of cell type names are even more sparse, with FlowCL being the only published method for labeling cell types based on their surface markers^41^. FlowCL similarly relies on the CL as a reference but lacks other features of this tool such as marker name standardization and species specification that offer additional flexibility. As far as we are aware, a tool that couples both the assignment of marker descriptions and descriptive cell type names has not been published. Notably, CytoPheno also encompasses all three parts into a graphical user interface (GUI). The GUI was included because many researchers find software that includes a user interface more useful and accessible than those without, particularly in the field of flow cytometry^33,42–44^. Feedback from various researchers was factored into the design of this interface and the tool is publicly available on GitHub.

## Methods

### Curation of Benchmark Datasets

All three parts of the CytoPheno tool were tested with expression data that contained cell clusters with delineated marker descriptions and descriptive cell type names (Table 1). Three benchmark datasets that satisfied these criteria and had distinct characteristics were used to illustrate the flexibility of the tool. The first benchmark set was mouse (strain C57BL/6J) bone marrow mass cytometry data from Samusik et. al (referred to as the ‘Samusik’ dataset)^45^. Only one sample, sample 1, was used from this dataset. The manually gated Samusik data was downloaded from the HDCytoData R package (FlowRepository ID: FR-FCM-ZZPH)^46^. The second benchmark was human PBMC mass cytometry data from Kimmey et. al (referred to as the ‘Kimmey’ dataset)^47^. This data is from a single donor and the 5-hour phorbol 12-myristate 13-acetate (PMA)/ionomycin stimulation set was used. The clustered Kimmey data was downloaded directly from FlowRepository (FlowRepository ID: FR-FCM-ZYR5)^48^. The third dataset was an in-house generated human PBMC spectral flow cytometry dataset (referred to as the ‘spectral’ dataset). This data included samples from seven healthy adult peripheral blood donors. The data was manually gated to get the specific cell type populations (Figure S1). Additional information about the sample preparation and the antibody panel can be found in the supplement (Table S1). This data is currently unpublished.

All datasets were transformed using an arcsinh function. The cofactors were set to 5 for the mass cytometry data and 6000 for the spectral data^49,50^. ‘Unassigned’ cells were removed from the Samusik and spectral data, as were markers that were used for preprocessing and markers that were not in the manual gating scheme at all. Similarly, markers that were not used in any marker definitions for the Kimmey dataset were removed. Since CD45^+^ is typically used for preprocessing, it was inconsistently used to define cell types within the benchmark datasets and so was excluded from all analysis. Heatmaps and histograms depicting the median and spread of the expression data for the included markers were made and examined for each dataset (Figures S2-S4, S5A, S6A, S7A).

**Table 1.** Datasets used to benchmark each part of the CytoPheno tool.

| Part | Benchmark | Type | Species | Platform | Sample type | # Markers used for phenotyping | # Cell types (original) | # Cell types used for testing (Part 3) |
| --- | --- | --- | --- | --- | --- | --- | --- | --- |
| 1-3 | Samusik | Expression values | Mouse | Mass cytometry | Bone marrow | 28 | 24 | 23 |
|  | Kimmey | Expression values | Human | Mass cytometry | PBMC | 11 | 9 | 7 |
|  | Spectral | Expression values | Human | Spectral flow cytometry | PBMC | 8 | 8 | 8 |
| 2 | Human marker set | Marker names | Human | Flow and mass cytometry | NA |  |  |  |
|  | Mouse marker set | Marker names | Mouse | Flow and mass cytometry | NA |  |  |  |
| 3 | OMIP 54 | Marker definitions | Mouse | Mass cytometry | Brain, spleen, bone marrow | 12 | 9 | 8 |
|  | OMIP 63 | Marker definitions | Human | Flow cytometry | PBMC | 23 | 28 | 23 |
|  | OMIP 78 | Marker definitions | Human | Flow cytometry | PBMC | 24 | 25 | 23 |

In addition to using the full expression data benchmark sets, the third part of CytoPheno – matching marker definitions to cell type names – was tested with three Optimized Multicolor Immunofluorescence Panels (OMIPs). These OMIPs were chosen to reflect different study designs including species, markers, and cell types. OMIP 54 was designed for studies using mouse brain, spleen, and bone marrow samples on the mass cytometer, while OMIPs 63 and 78 were created for human PBMC flow cytometry studies^12,51,52^. The cell types included for benchmarking for OMIPs 54 and 78 were those that were numbered in the respective manual gating strategies (Figure 1 in both OMIPs)^12,52^. OMIP 54 also included several activation states for certain cell types that were excluded due to naming ambiguity. Since OMIP 63 did not include numbered cell types in the manual gating figure, the included cell types were instead taken from a cell type definition table included in the manuscript (Table 3 in the OMIP)^51^. For each OMIP the manual gating figure, in combination with textual descriptions, was used to determine the markers that describe the cell types.

Descriptive cell type names within the Samusik, Kimmey, spectral, OMIP 54, OMIP 63, and OMIP 78 benchmarks were matched to the closest corresponding CL term, which served as the ground truth (Tables S2-S7 respectively). ‘Closest’ was determined by comparing the CL descriptive name and marker definitions with those of the benchmark datasets and finding the most specific corresponding match. While the Samusik dataset did include CL terms from an older version of the CL, some terms in the current CL were determined to be more accurate matches, thus these terms were used instead. When considering all the benchmark sets, there were some cases where two CL terms matched equally well to a benchmark cell type. In these instances, both terms were considered the ground truth. In more ambiguous instances where three or more terms matched equally well, that cell type was excluded completely from the benchmark set. Other cell types were excluded from testing when there was no CL term that corresponded to the cell type described in the benchmark data.

To further test the marker standardization part (Part 2) of the tool, additional testing data was used to encompass more markers and get more representative examples of naming conventions researchers use in experiments. The human test dataset was made up of user reported markers from 15 flow and mass cytometry studies in the Immunology Database and analysis Portal (ImmPort) (Table S8) that were initially released between December 19^th^, 2022 and December 15^th^, 2023 (data release versions 46–50)^53,54^. The marker names were extracted from metadata in the ‘reagent’ file (specifically the ‘analyte_reported’ column), with other pertinent study information included in the ‘study’, ‘subject’, and ‘biosample’ metadata files. Markers that were less than three characters and markers named as the internal ImmPort analyte identifier were excluded. Special characters that were not able to be read accurately in the R Studio environment were removed. Entries were split by semicolons, since this character was used to put multiple markers together (note that other characters were sometimes used for this purpose, but those characters also had incongruent uses so they could not be universal separators). The final number of unique markers was 176, with ‘unique’ defined as being capitalization sensitive and including punctuation and spaces. After all characters were capitalized and dashes, underscores, periods, and spaces were removed, the number of markers was 158.

Since ImmPort studies from this timeframe only encompassed human samples, mouse studies in FlowRepository were used to test the workflow and evaluate its consistency across species. Conventional flow, spectral flow, and mass cytometry studies in FlowRepository that included the word ‘murine’ in the listed experiment name, with an ‘updated’ date between January 2021 and December 2023 (searched on January 15^th^, 2024) were included in the test set. For each study a Flow Cytometry Standard (FCS) file was randomly chosen and downloaded. The flowCore package was used to open the FCS file in R and the markernames function was used to extract the marker names^55^. The FCS files for 2 studies did not have protein names listed in that manner and so were removed. Ultimately, 10 studies (Table S8) remained in the murine test set, encompassing 199 unique marker names. ‘Unique’ was defined in the same manner as in the human test set. After all characters were capitalized and dashes, underscores, periods, and spaces were removed, the number of markers was 197.

### Designation of Marker Descriptions (Part 1)

An algorithm that requires post-clustered expression data and concludes with each phenotyping marker being defined as positive, negative, or left without a specific designation (null) was produced using multiple developmental datasets for testing.

The algorithm first calculated the median expression value of every cluster for every inputted marker (labeled the ‘cluster of interest’). Then, the median of the combined cells from all other clusters for that same marker was extracted (called the ‘reference clusters’). To account for cases where a marker may be completely positive or completely negative across clusters, threshold values for these medians and the standard deviation of the entire dataset were used. As with the algorithm itself, all default values were determined based on developmental data. Specifically, a marker was designated as null across clusters if all cluster medians were below a certain value (default = 0.7), the minimum median across those clusters was below another value (default = 0.2) and the overall standard deviation was below a third value (default = 0.5). A marker was designated as positive across clusters if all cluster medians were above a certain value (default = 0.2), the maximum median across those clusters was above another value (default = 1), and the overall standard deviation was below a final value (default = 0.7).

After accounting for cases where a marker was completely negative or positive across clusters, other markers were designated as negative, positive, or null per cluster based on the median difference between the cluster of interest and the reference clusters (Equation 1). After the median differences were calculated, the differences were rescaled for each marker with a min-max normalization between −1 and 1 (Equation 2). A marker was designated as positive if the scaled value was above a specific threshold (default = 0.25). Likewise, a marker was assigned negative if the scaled value was below a specified threshold (default = −0.75). Values between the positive and negative cutoffs were labeled as null, as there was low confidence in determining if they were negative or positive. The Samusik, Kimmey, and spectral benchmark datasets were used to test this part of the CytoPheno tool. All original cell types were used in this analysis. The ground truth marker definitions were compared to the algorithm produced marker definitions through a binary classification method (Table S9). Accuracy and the true positive rate were assessed. Please see the supplement for additional details regarding the binary classification method.

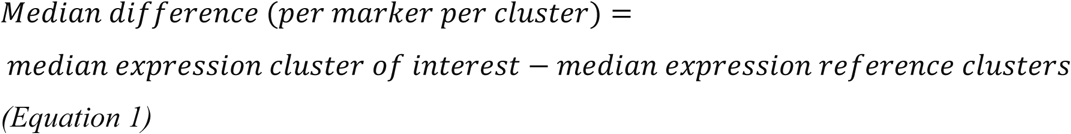

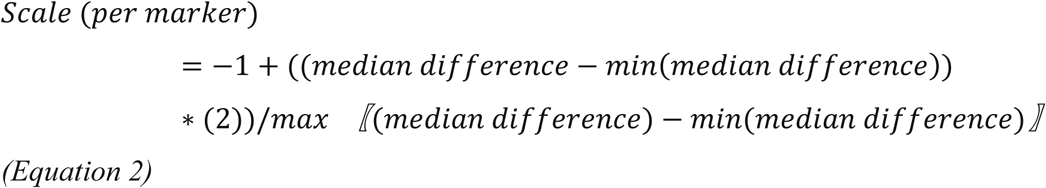

### Standardization of Marker Names (Part 2)

A multistep workflow that begins with user-inputted protein markers and ends with the corresponding PRO or GO terms was created^34–36^. Linking inputted marker names to PRO or GO terms was necessary to ultimately query against the CL in Part 3^37^. This workflow was developed with data downloaded from the ImmPort Repository^53,54^. Specifically, metadata encompassing all flow and mass cytometry studies with initial release dates including and prior to September 2^nd^, 2022 (<= data release version 45) was used. This developmental data was extracted in the same manner as that of the human benchmarking dataset. This set contained markers from 127 studies from 4 species: human (105 studies), mouse (18 studies), rhesus monkey (3 studies), and night monkey (1 study) studies (Table S10). The human set initially contained 619 unique (capitalization and punctuation sensitive) markers, while the mouse set had 203 markers, the rhesus monkey set 55 markers, and the night monkey set 15 markers. After all characters were capitalized and dashes, underscores, periods, and spaces were removed, the number of remaining markers was 563 in the human set, 185 in the mouse set, 55 in the rhesus monkey set, and 15 in the night monkey set.

These markers were subsequently examined to determine the best methods for standardizing the marker names. While SPARQL Protocol and RDF Query Language (SPARQL) queries directly to Protein Ontology was the primary approach, these queries sometimes failed due to a widespread lack of consistent protein naming conventions. As a result, a multistep process that uses several naming references was created (Figure 2). Importantly, the developmental data was also used to manually create the suggestion lists used in the workflow. Please see the supplement for details regarding the full development of this workflow.

**Figure 2:**
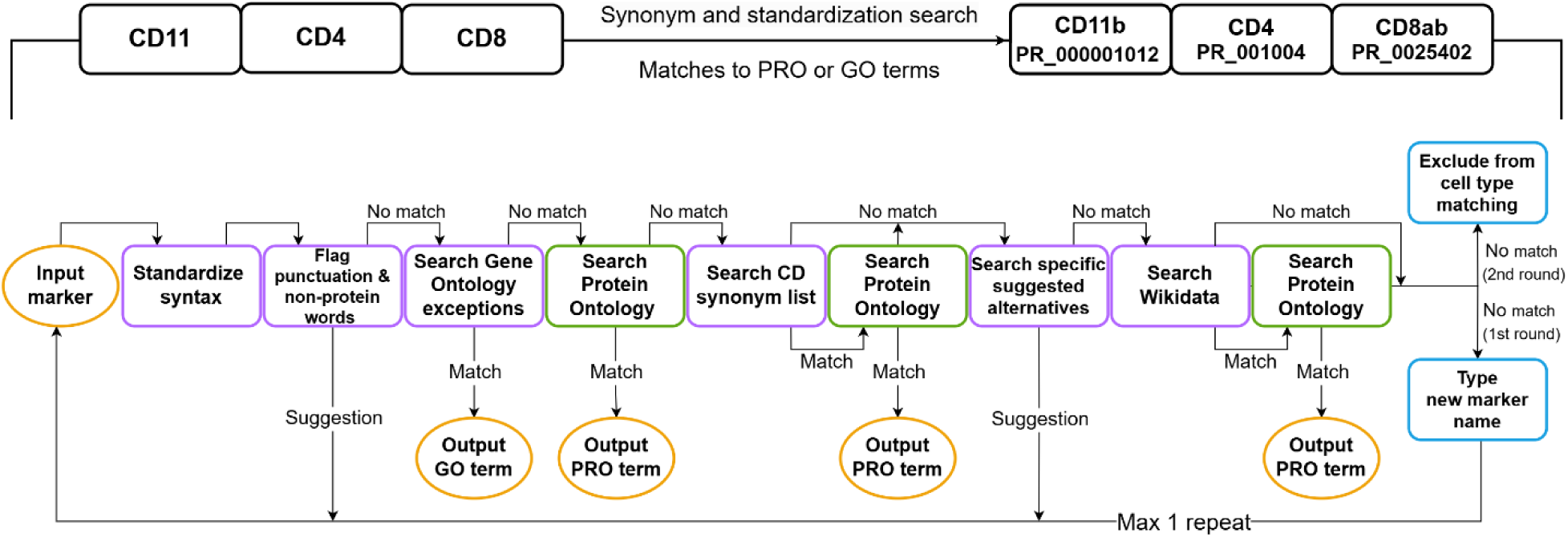
Part 2 workflow from initial marker name to final standardized PRO or GO linked term.

Tests with the benchmark datasets (see ‘Curation of Benchmark Datasets’) were conducted to see how well this workflow would standardize marker names with new data. Both marker sets were put through the created workflow and the results were recorded.

### Descriptive Naming of Cell Types (Part 3)

Markers standardized to PRO or GO terms, along with each marker’s categorical expression level identifier (i.e. positive and negative) were used to match the marker descriptors to cell type names for each cluster. SPARQL was used to directly query CL and PCL. The human-readable qualifiers were converted to equivalent Relations Ontology terms, which are used in CL class relations definitions^56^. Specifically, positive reverted to ‘has plasma membrane part’ (RO_0002104) and ‘has part’ (BFO_0000051), high to ‘has high plasma membrane amount’ (RO_0015015), and low to ‘has low plasma membrane amount’ (RO_0015016). Negative corresponded to the CL term ‘lacks plasma membrane part’ (CL_4030046). The combination of the PRO or GO terms with these qualifiers was then used to match cell type classes, which have structured definitions that specify the presence or absence of proteins. Since CL is a hierarchy, each cell type definition encompasses both that specific individual cell type and all the cell type definitions in the parent lineage (superclasses).

For development and testing purposes, a marker match between a marker within the inputted marker definition and a marker within the CL marker definition was defined as any time the same PRO or GO term had exactly the same negative or positive qualifier. Positive markers also matched to high and low markers. For high markers a match was defined as anything that was also high or positive, but not low. For low markers a match was defined as anything that was also low or positive, but not high. A contradiction was defined as any instance when the same input and CL marker had a qualifier that was not a match. For example, an input of ‘CD4^+^’ would be a match to ‘CD4^low^’ but a contradiction to ‘CD4^-^’.

Although over 200 different PRO or GO terms are currently used to define cell types in the CL, not all potential marker names are included in this ontology. To account for this, inputted marker names that were not in the CL were excluded entirely from the scoring process. This included all markers that could not be matched to a PRO or GO term, as well as those linked to a PRO or GO term that was not in the CL.

All CL cell types that contained at least 1 match with the inputted marker description were scored and ranked by that score to determine which cell types were a more optimal match. A modified Jaccard similarity equation was used to score the results (Equation 3).

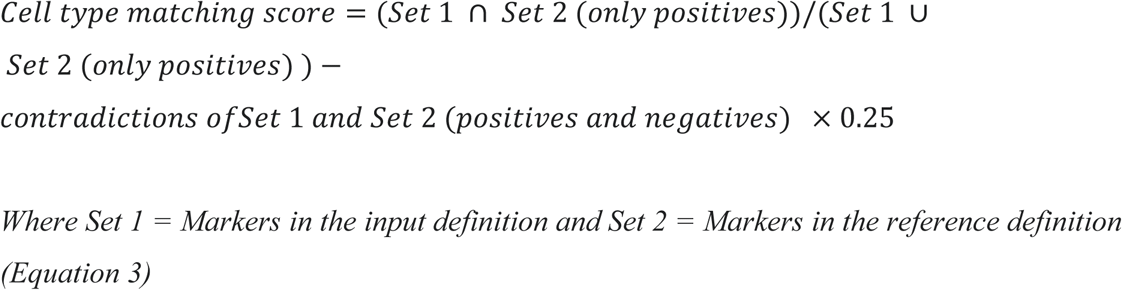

Only positive marker designations were used to define the intersection and union because researchers tend to explicitly define cell types with a larger proportion of positive marker designations compared to negative designations. Both positive and negative contradicted markers were used since all contradictions should be considered a concern and penalized as such. The Samusik, Kimmey, and spectral expression datasets, as well as the OMIP 54, OMIP 63, OMIP 78, were used to test the performance of this ontology cell type matching method and scoring system by comparing results to the ground truth cell type names. Performance was assessed by score rank, with lower numbers indicating better matches. In other words, a ground truth cell type name that had the third highest score would be ranked as a ‘3’. The methods top-3 accuracy and top-5 accuracy were used to assess the results in a categorical manner, as has been done previously^57^. Top-3 encompassed any rank in the top three and top-5 any rank within the top five. The January 29^th^, 2025 version of the Cell Ontology was used to attain all benchmark results.

### Formation and Usage of the Graphical User Interface

The Shiny R package was used to develop the graphical user interface (GUI)^58^. Additional R packages were used to add features to the application, such as input validators (shinyvalidate), loading icons (shinybusy), data tables (DT), and the ability to enable and disable elements (shinyjs)^59–62^. The application was designed to incorporate the entire process from post-clustering expression data to cell type names in an easy-to-use manner.

Specifically, each of the three broad parts are broken down into substeps. Part 1 has three steps: 1) choosing the markers, 2) choosing the median difference parameters, and 3) overriding the results of the median difference parameters. Part 2 has two steps: 4) matching to PRO or GO terms (and reinputting marker names if necessary) and 5) choosing which terms to accept. Part 3 has one step: 6) matching to descriptive cell type names.

Notably, each substep outputs tables or figures that depict the data and results. Visualizations are in the form of heatmaps (heatmaply) and density plots (ggplot2)^63,64^. Most plots and tables are downloadable for off-application usage. Additionally, each step provides flexibility, allowing users the option to change parameters or override results. Users can initially indicate if any markers were used for pre-gating, if the expression data should be arcsinh transformed, if the data should be subsampled, and if a seed number should be specified. In Part 1, users can also remove any markers from analysis, change all median difference parameters, and override any specific results. For Part 2, species can be specified to allow both species specific PRO and CL terms to be selected for. Additionally, unmatched marker names can be edited and any markers can be removed from subsequent analysis. For Part 3, the cell type ontology (CL, PCL, or both) and the number of cell types returned per input cluster can be specified.

Users can choose to bypass Part 1 (substeps 1– 3) by using their own marker definition assignment method and begin at Part 2 in the tool. Starting at Part 2 requires inputting marker definitions rather than expression data but follows all the same subsequent steps. When inputting marker definitions, low and high expression can also be denoted. The interpretation of these qualifiers can also be specified, since the interpretation of both low and high can be ambiguous and disjointed amongst researchers. Specifically, low can be indicated as either matching to low alone, matching to low and positive qualifiers, or matching to low and negative qualifiers within the CL. High can similarly be specified as matching to high alone or matching to high and positive qualifiers within the CL.

This work is focused on using non-uploaded (default) references, which include both curated suggestion data sets and data extracted from a variety of online sources (including CL, PRO, GO, Wikidata, UniProt, InterPro, Protein Data Bank, and Human Cell Differentiation Molecules’ CD list)^34–37,65–70^. However, to account for more potential usage cases, the application also includes the option to use uploaded references in place of default references. This reference should be a marker-cell type table that contains cell type names and associated marker definitions. When selecting this option, Part 2 is ignored and only the uploaded reference table is used for Part 3.

## Results

The CytoPheno tool was tested with multiple benchmark datasets (Table 1) that demonstrated the performance of each part of the tool.

### Assigned Marker Descriptions Reflect Ground Truth Definitions (Part 1)

In Part 1, unidentified clusters are assigned marker definitions (positives, negatives, or nulls) based on marker expression values. Three cytometry datasets, two mass cytometry (Samusik and Kimmey) and one spectral flow cytometry (spectral), that contained cell clusters (from unsupervised clustering or manual gating) and the corresponding marker definitions were used. This encompassed 41 different cell types. The outputted marker definitions (Figure S5B, S6B, S7B) from the Part 1 algorithm were then compared to the ground truth definitions through binary classification methods (Figure 3). For the Samusik, Kimmey, and spectral sets, the accuracy values were 0.888, 0.862, and 0.809 respectively. The true positive rates for those datasets were 0.849, 0.875, and 0.789 respectively.

**Figure 3.**
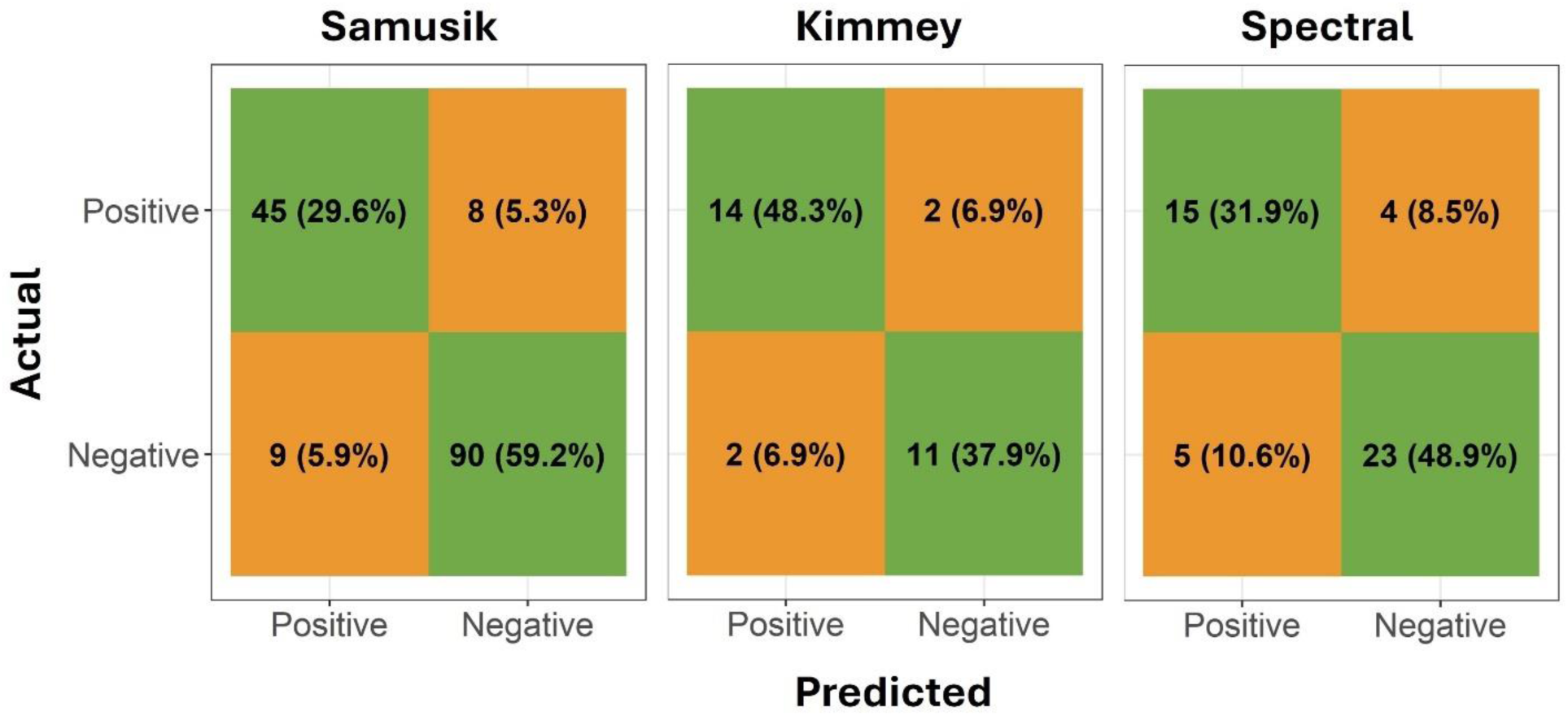
A 2×2 confusion matrix depicting the number (percentage) of actual and predicted markers categorized as positive or negative for each of the three (Samusik, Kimmey, and spectral) benchmark datasets.

Importantly, when both actual positives and actual negatives were misclassified, it was almost always assigned a ‘null’ value. Between the three datasets, there were no instances of a true negative being misclassified as a positive. There was a single instance when an actual positive was misclassified as a negative. This occurred in the Kimmey dataset, where researchers defined non-classical monocytes as CD14^low^ and CD16^+^. ‘Low’ was considered ‘positive’ during benchmarking, although in this case there is some ambiguity as to the meaning of ‘low’ since heatmaps and density plots of the underlying expression data indicated negative expression (Figures S3 and S6A).

### Standardized Marker Names Match to Ontology Terms (Part 2)

In Part 2 of CytoPheno, marker names are standardized. Specifically, this part aims to translate user-inputted marker names to the standardized names and identification terms from PRO and GO. Since the marker names from the other benchmark datasets were already published with mostly standard naming conventions, this part of the tool was developed and tested with additional data from ImmPort and FlowRepository^48,53,54^. ImmPort contains metadata with original marker names as inputted by users in their FCS files, while the marker names can be directly extracted from FCS files in FlowRepository. The developmental data was used to determine which protein references were useful, the order of those references in the multistep workflow, and which suggestions to include in the curated user-suggestion lists within the workflow (Figure 2).

**Table 2.** Results of the benchmark testing for Part 2 (Standardization of Marker Names). Testing data was extracted from ImmPort (human test set) and FlowRepository (mouse test set). Each match step is listed, along with the number and percentage of markers that matched from the human and mouse test sets. ‘Matches’ refer to match steps that returned a GO or PRO term without human edits. ‘Suggestions’ refer to steps that returned a suggestion of how to edit the marker name to the user. ‘Unmatched markers’ were markers that did not match to a PRO or GO term and did not receive any suggested edits.

|  | **Match Step** | **Human test set,**<br>**N (%)** | **Mouse test set,**<br>**N (%)** |
| --- | --- | --- | --- |
| **Matches** | GO exceptions | 9 (5.11%) | 7 (3.52%) |
|  | Directly to PRO | 105 (59.66%) | 65 (32.66%) |
|  | CD synonym list | 2 (1.14%) | 1 (0.50%) |
|  | Wikidata | 1 (0.57%) | 1 (0.50%) |
| **Suggestions** | Specific protein suggestions | 5 (2.84%) | 4 (2%) |
|  | Non-protein word suggestions | 11 (6.25%) | 111 (55.78%) |
|  | Punctuation suggestions | 38 (21.59%) | 5 (2.51%) |
| **Unmatched markers** |  | 5 (2.84%) | 5 (2.5%) |

Two benchmark datasets, one with marker names from human studies and the other with marker names from mouse studies were used for performance assessment (Table S8). Assessments were based on whether the marker name matched to at least one PRO or GO term (Table 2). Whether or not this matched term was the ‘correct’ match was impossible to assess without knowing the user’s intentions and so could not be evaluated.

For the human set, 117 (66.48%) markers were able to be matched to terms without requiring additional user input (Table 2). 54 (30.68%) markers were not able to initially be matched but a specific suggestion was provided to the user that could be followed to correct the marker name so it would successfully match during a second run through the workflow. Only 5 (2.84%) markers were neither able to be matched nor were they encompassed within the suggestion lists. For the mouse set, 74 (37.19%) markers were matched directly to a term and 120 (60.30%) markers were not matched but returned a suggestion to the user. Numerous markers in the mouse set contained metal tags as part of the ‘marker names’ (e.g. 151EuCD25), accounting for the large percentage of markers that required a suggestion, namely that a metal tag is included as part of the name and that it should be removed. Still, only 5 (2.5%) markers were not able to be matched at all.

The marker names from the benchmarks introduced in Part 1 (Samusik, Kimmey, and spectral datasets) as well as data from three Optimized Multicolor Immunofluorescence Panels (OMIPs) were also matched to PRO or GO terms to prepare for cell type naming within the CL (Tables S11-S16). Within these 6 benchmarks, there were 106 markers, 58 which represented unique inputted marker names. Only one marker (120G8) could not be matched to any GO or PRO term, while two markers (TCR Vα24JαQ and TCR Vα7.2) were not specifically represented within PRO, but still could be matched to the more general TCR protein complex.

### Matched Cell Types Reflect Ground Truth Names (Part 3)

CytoPheno ends with Part 3, where the cell clusters are given descriptive cell type names through the Cell Ontology and Provisional Cell Ontology. Six benchmark datasets (Samusik, Kimmey, spectral, OMIP 54, OMIP 63, and OMIP 78) were used for testing. For the Samusik, Kimmey, and spectral sets, the results of Parts 1 and 2 were inputted into Part 3. For the OMIPs, the manual gating and corresponding textual definitions were used to identify cell types and their marker descriptions. The marker names were standardized in Part 2, with the results similarly inputted into Part 3.

The six benchmark datasets encompassed numerous cell types, many of which were unique. Specifically, 103 cell types were defined in the benchmark sets. Of these 103 initial cell types, only 11 could not be matched to a corresponding CL cell type. Eight cell type definitions were too broad and equally matched with too many cell types in the CL to assess (>2 cell types). Only 3 cell types (atypical memory B cells, CD16^low^ CD56^+^ natural killer cells, and CD16^+^ CD56^-^ natural killer cells) were specific and still had no equivalent cell type definition within the CL or PCL. Additionally, there were 6 cell types that equally matched to 2 cell types, which was included in the analysis (the score rank between the two was averaged for assessment purposes). As a result, there were 92 cell type definitions included in the benchmark analysis, which matched to 98 total CL terms. While the 92 cell type definitions contained many repeated markers, no two cell type definitions were exactly the same. Additionally, while some of the benchmark cell types did correspond to the same CL terms, 67 different CL terms were still represented within the 6 benchmarks.

**Table 3.**
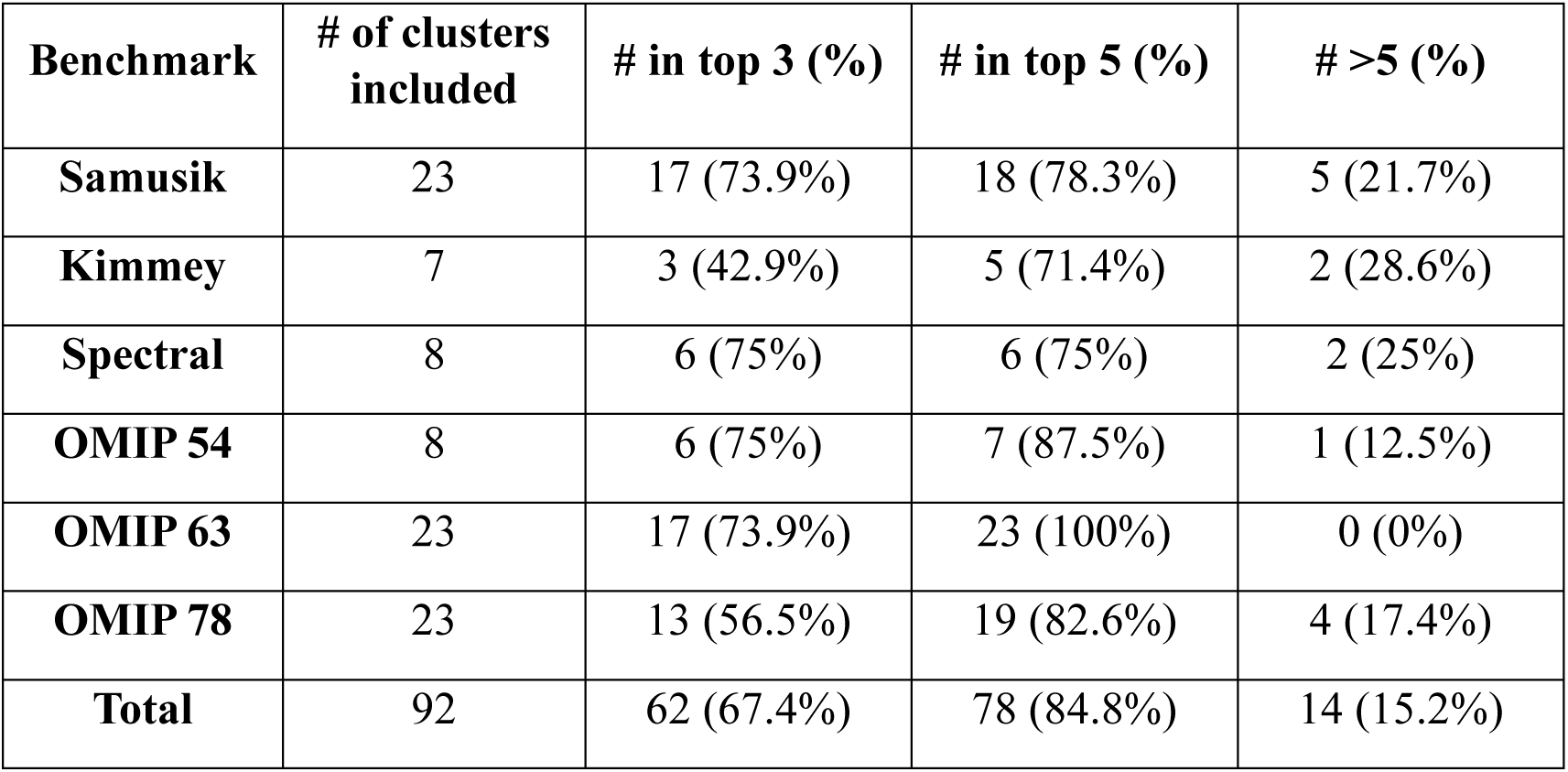
Results of the benchmark testing for Part 3 (Descriptive Naming of Cell Types). Each benchmark is listed, along with the number of clusters that were included in the ontology matching search. The number and percentage of clusters where the ground truth cell type was within the top-3 and top-5 ranked matches are listed, as well as the number and percentage of clusters that were not within the top matches.

Once the ground truth CL terms were determined, the benchmark marker descriptions were matched to those within the CL and PCL. The quality of the match was quantified through a score (Equation 3). The matches were ranked by highest to lowest score and the ranking position of the ground truth cell type was recorded. Ranks were considered categorically, using the top-3 and top-5 accuracy methods (Table 3). Of the 92 potential cell types, 62 (67.4%) ranked within the top 3 scores and 78 (84.8%) were ranked within the top 5 scores.

Only 14 (15.2%) were ranked outside of the top 5 scored cell types. In most cases, the poorly ranked matches were due to markers used within the ground truth definition that were not delineated in the corresponding CL cell type. There were also some cases where the ground truth marker was defined as negative and the CL cell type had that same marker defined as low, or vice versa. Additionally, the results do indicate that the OMIPs performed slightly better overall than the Samusik and Kimmey datasets, which is unsurprising since the latter expression values went through every part of the tool while the former marker definitions were only processed through Parts 2 and 3.

### Graphical User Interface to Enhance Tool Usability

Finally, a graphical user interface was created in R Shiny that encompasses all three parts of CytoPheno (Figure 4). This application was tested by both wet-lab biologists and computational scientists, with user feedback incorporated into its design. The GUI allows non-programmers to easily and freely use the tool by going through all or some of the three main parts and six substeps. The application allows researchers to visualize, make edits, and download their results.

**Figure 4.**
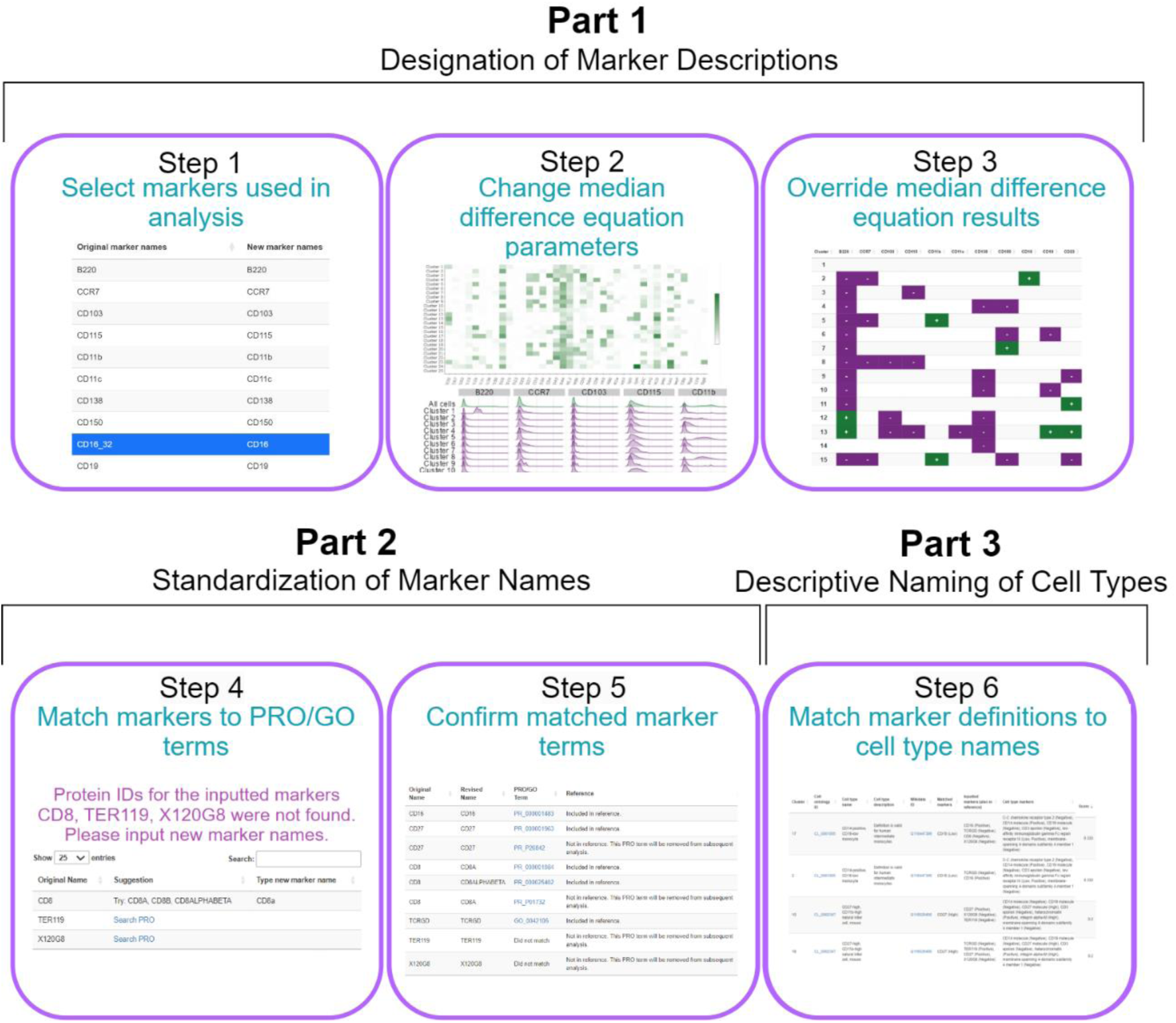
Simplified depiction of the three parts and six substeps that form the R shiny GUI. Each step represents a separate section the GUI takes the user through. The boxes contain a brief description and an example screenshot from that part of the application. The screenshots depict certain sections, but not the entirety, of the user interface.

Specifically, the final output of Part 1 includes the results of the median difference equation, with each marker specified as being positive, negative, or null per cluster. Various textual outputs of these results can be downloaded as CSV files. Additionally, a binary heatmap is created that depicts this information in plot form. Other plots include various heatmaps and a density plot depicting the underlying marker expression values. All plots can easily be downloaded for off-application usage.

For Part 2, the final output is a data frame with all inputted marker names, revised marker names (if applicable), and the corresponding PRO or GO terms, with direct links to the online PRO or GO entries. These links allow the user to easily check if the matched standardized marker names correspond with their intentions. Specifically, multiple PRO terms can match, since many markers have both species distinct and non-species-specific terms. Additionally, some inputted marker names are too ambiguous and may be listed as a synonym of multiple terms. The output also informs the user if the marker is encompassed in one or more marker definitions within the CL.

For Part 3, the final results are displayed within a data frame that includes columns specifying the input cluster, the matched CL cell type name, the CL identification term, a description of the matched cell type, and a Wikidata identification term. Both the CL and Wikidata terms are clickable and lead to their respective web pages. Additionally, the markers that are in the input cluster, markers that are present in the matched CL cell type, markers that are matched between the input cluster and CL cell type, and markers that are contradicted between the input cluster and CL type are included. Finally, the output includes three metrics for evaluating the match. These include the percentage of matched markers over the total markers in the CL cell type, the percentage of matched markers over the total markers in the input, and the matching score (Equation 3).

Both the application itself and an extensive user guide for the application are available on GitHub (https://github.com/AndorfLab/CytoPheno).

## Discussion

Recent advances in flow and mass cytometry have increased data complexity, thereby increasing the necessity for computational tools to interpret the data^71^. The weaknesses of a manual approach to flow and mass cytometry data analysis is well documented^3–7^. Principally, manual analysis of large datasets is laborious, inefficient, subjective, and hard to reproduce^71^. Automated gating strategies have addressed these drawbacks, however post-gating tools that aim to explicitly characterize cells by their marker expression patterns and assign descriptive cell type names are lacking^21^. The cytometry phenotyping tool presented here fills this gap through a series of steps that aim to connect unnamed cell clusters to marker descriptors, standardized marker names, and standardized cell type names.

While CytoPheno does automate marker definitions and cell type naming of unnamed clusters, a researcher should still provide some manual analysis to ensure that the step-by-step results are consistent with the underlying data. In particular, the GUI promotes this by walking the user clearly and concisely through the three parts and their six substeps. The interface plainly shows results that can then be overridden if need be. Heatmaps and density plots are produced through the tool to aid in this endeavor. Secondly, while all three parts of the tool can be easily used in combination, this is not a necessary requirement. The user can skip Part 1 and input their own marker descriptions that were either manually derived or outputted by another algorithm. Equally, Parts 2 and 3 can be ignored if the user chooses only to output the results of Part 1. Alternatively, the incorporated default references that encompass Parts 2 and 3 can be replaced by an external reference. A user can upload a marker-cell type reference table, such as the ones used by some semi-supervised methods^25,31^. Another option is to use a standardized panel for the reference, such as one that has been developed and used across experiments in large consortia^5^. Using an uploaded table may better incorporate the cell type definitions a user is seeking and offers a speed advantage, but could also retain a level of subjectivity and is limited to provided cell types.

Benchmark datasets were used to illustrate the tool’s ability to assist researchers in the cell type annotations process. Importantly, these well-defined datasets were chosen to include both human and mouse data, as well as a variety of data designs to illustrate the breadth of this tool. For Part 1, the tool was created to maximize the predictive power of true positives (markers with a positive expression designation). This strategy was pursued because positive expression of markers is oftentimes the primary method of distinguishing different cell types and so analysts tend to explicitly state more positive designations than negatives, while many markers are left unspecified. Therefore, while broad measures of performance are still useful, the true positive rate (sensitivity) was given primary weight in development and assessment of the algorithm. Indeed, benchmark sets achieved high true positive rates, indicating success in this regard. Human inputted marker names were taken from the FlowRepository and ImmPort repositories to illustrate the performance of Part 2. The results indicate the multistep workflow for getting PRO and GO terms is thorough. When PRO or GO terms were not able to be matched, it was typically because of bad naming conventions by the user (i.e. metal or fluorochrome was still attached to the protein). However, most of the time CytoPheno was able to flag why the protein was unable to match and prompted the user to make specific revisions. Each of the remaining markers that were not able to be matched or given a suggestion for matching were examined independently to determine the cause. Of the 10 unmatched markers (<3% of the inputted markers), 3 were in a format that did not represent an actual, singular protein marker (CD8_IgD, I-A_I-E, phosphatidylserine), 2 represented a protein in PRO that had an insufficient name (arginase, siglec-8 ligand binding), and 5 represented antibodies whose names were not in PRO at the time of this analysis (ivCD45_2, Siglec-H, GL7, pSTAT5, TCR Va7.2). To improve future performance of the tool, ‘phosphatidylserine’ was added to the non-protein word list and ‘arginase’ and ‘siglec-8 ligand binding’ was added to the specific suggestion list (which now suggests changing to ARG1 and SIGLEC8 respectively).

Six benchmark datasets were used to demonstrate Part 3 of the tool, the annotation of descriptive cell type names. These datasets were used to represent a variety of data types and show how frequently different naming conventions and marker definitions are used by researchers. However, the majority (89.3%) of cell types in the benchmarks still were represented within the CL and used for assessment. Cell types that were defined in the benchmarks very broadly (matched equally well to more than two CL cell type terms) were excluded from the benchmark assessment because it was hard to determine a correct match. When a benchmark cell type equally matched to two CL terms that cell type was included, with the final rank taken as the average of the rank of both CL terms.

Only three cell types from the benchmark datasets did not have an equivalent term in the CL or PCL, indicating the breadth of cell types represented in these resources. Due to the large amount of cell types represented in the CL, some of which only differ from another CL term by a single surface marker or in other cases no markers at all, and the large amount of potential cell type marker definitions in the literature, it was expected that conventional accuracy (top-1) would not be a sufficient output or metric for the user. Instead, the tool returns multiple matches, ranked by the scoring method. Therefore, performance was assessed through the top-3 accuracy and top-5 accuracy methods, which has similarly been used in scRNA-seq CL annotation methods^57^. The top-3 and top-5 accuracy methods resulted in accuracies of 67.4% and 84.8% respectively. This suggests researchers should examine multiple results and interpret them in conjunction with their underlying data when using CytoPheno. Ultimately, as with most automated tools, the researcher’s expert knowledge of their experimental design should guide the final assignment of cell type names.

While higher level cell types may be fairly well-defined, new markers and more specific cell types are still being discovered, making the process of phenotyping increasingly complex data more time-consuming, biased, and error prone^72^. The tool’s strength lies in its ability to combat these obstacles by harnessing the power of curated structured resources and ontologies to provide large amounts of standardized marker names, cell type names, and cell type definitions. Currently, researchers use a variety of naming conventions to refer to the same protein marker or cell type^73^. Conversely, some cell type names are consistently used by researchers across institutions, but these same names may have different marker definitions. Benchmark data used to test the tool illustrates both types of ambiguity (Tables S2-S7). Overall, the uncertainty of how cell types are named and defined can have a meaningful impact on how others interpret reported results, emphasizing the need for better naming practices in the field. Importantly, larger meta-analyses, cytometry data integration, and multi-omics integration techniques could be aided through standardized naming conventions.

The large amount of biomedical knowledge openly available to researchers offers promising potential for clarity, but only if that data can be easily accessed and is consistent across platforms. The usage of biomedical ontologies that make up the Open Biological and Biomedical Ontology (OBO) Foundry offers a solution^74,75^. OBO follow standard principles that allow users to harness the power of a controlled vocabulary that structures entities by biological class definitions and their relations. Primarily, CytoPheno relies on PRO and CL. Notably, the breadth of knowledge from the CL has previously been leveraged to aid in establishing cell type annotations in various scRNA-seq applications^57,76–78^. Since gene expression does not always equate to protein expression, these tools themselves are not optimal recourses for cytometry analysis but their existence does illustrate the increasing role ontologies play in providing standardized naming conventions in single-cell analysis^79,80^. Annotating both transcriptomics and proteomics data with ontological terms may also benefit future multi-omics data analysis through interoperable naming conventions.

Several considerations should be given when using CytoPheno and interpreting the results. First, the tool was optimized for immune cell types, as they are typically classified according to expressed cell surface proteins^72^. The CL does contain non-immune cell types that may be classified by other features such as morphology, but the tool is not currently capable of querying for such features. A second consideration is the CL’s current focus on cell types in a healthy homeostatic state, rather than in disease states^81^. Additional resources, such as disease specific ontologies, could be integrated into the tool in future work to provide additional usefulness to research focused on various pathological states. Thirdly, while the CL officially represents both mammalians and non-mammalian vertebrates, there is most likely bias towards human and mouse cell types because they are much more studied and described compared to those of other species^37^.

CytoPheno has several limitations. Although Part 1 allows the user to optionally change cutoff values to better suit individual data needs, Parts 2 and 3 rely on resources that cannot easily be altered. While users can submit changes and updates to OBO, this tool’s reliance on those resources will not allow for immediate amendments. This limitation is mitigated by the wide breadth of the ontologies, as the CL and PCL each contain about 3,000 cell type terms, while PRO currently includes over 250,000 protein terms. Additionally, because ontologies are continuously updated and the tool queries directly from a web-based linked server, it is expected that this tool will become more extensive and inclusive as additional term modifications or additions are made^81^. For example, both OMIPs that utilize well-studied surface marker characterizations and comprehensive studies that characterize cell surface protein expression per immune cell type offer additional cell type-marker combinations that can be integrated into the CL^82–84^.

Overall, CytoPheno offers users an efficient post-clustering approach for characterizing undefined cell types by both their marker descriptors and informative cell type names. Notably, the tool includes multiple parts that can be used in unison or separately, allows for different input and reference formats, and includes an easy-to-use GUI. These aspects promote the tool’s flexibility and accessibility, allowing researchers from a variety of backgrounds with differing study designs and aims to use the tool.

## Data Availability

The Samusik et al. and Kimmey et al. datasets analyzed in this paper were previously published and are available on FlowRepository, with repository identifications FR-FCM-ZZPH and FR-FCM-ZYR5 respectively^45,47,48^. The spectral dataset analyzed in this study is currently unpublished but will be made available in an open repository upon final manuscript publication. The human and mouse marker datasets used openly available data extracted from the ImmPort Repository and FlowRepository (See Tables S8 and S10 for all repository identification numbers)^48,53,54^. Data from the Optimized Multicolor Immunofluorescence Panels 54, 63, and 78 were taken directly from the respective manuscripts^12,51,52^.

## Code Availability

The R Shiny application created and used in this study is available on GitHub at https://github.com/AndorfLab/CytoPheno.

## Competing Interests

The authors declare that they have no conflict of interest.

## Ethics

The spectral data is classified as an “exempt” research project. All other data used in this study was repurposed, with ethical approvals secured by the original study authors.

## Funding

This work was supported in part by the National Institutes of Health (NIH) grant P30AR070549 (S.T., S.A.); a Trustee Award from the Cincinnati Children’s Research Foundation (CCRF) (S.A.); a CCRF Trustee Award (T.T.); and the Burroughs Wellcome Fund Next Gen Pregnancy Award #NGP10115 (T.T.). The purchase of the Cytek Aurora was funded by NIH grant S10OD025045. Additionally, T.T. is supported by the March of Dimes Prematurity Research Center Ohio Collaborative. The content is solely the responsibility of the authors and does not necessarily represent the official views of the NIH.

## Notes

### Competing Interest Statement

The authors have declared no competing interest.

