## Supplement for "CytoPheno: Automated descriptive cell type naming in flow and mass cytometry"

#### Spectral Data Sample Preparation and Cytometer Analysis

Peripheral blood was obtained from healthy, non-pregnant adult donors. All human samples used for this research were de-identified, with discarded clinical material and no patient or clinical information available for analysis. Whole blood from each donor was diluted 1:1 in 1x PBS to a final volume of 50 mL and 25 mL aliquots were layered on 12 mL Ficoll gradient. After gradient centrifugation (20 min, 2000 rpm), lymphocytes were isolated from the Ficoll interface and washed once in RPMI containing pen/strep and 10% FBS and directly stained for analysis on a 5-laser Cytek Aurora spectral flow cytometer.

Antibodies and isotype controls used for flow cytometric analysis are listed in Table S1. For surface staining, cells were stained for 30 min in RPMI containing pen/strep and 10% FBS. For intracellular staining, cells were fixed and permeabilized using the eBiosciences FOXP3 kit (eBiosciences). Analysis was performed on a Cytek Aurora. All flow cytometric data were acquired using equipment maintained by the Research Flow Cytometry Core in the Division of Rheumatology at Cincinnati Children's Hospital Medical Center.

#### Binary Classification

Binary classification was used to assess the performance of the Part 1 algorithm. Positive and negative marker designations were each considered their own condition, while null marker designations were only considered when a ground truth positive marker designation or a ground truth negative marker designation was labeled as null. Null values were only judged in this manner for a few reasons. One reason was that cases of actual nulls corresponding to predicted nulls could inflate accuracy numbers since there are many nulls that are in both the ground truth and are predicted as such. Additionally, instances where benchmark nulls were predicted as being either positive or negative were not considered since the Kimmey marker definitions and the Samusik and spectral manual gating data did not contain comprehensive cell type marker definitions but rather focused on the most 'important' or defining markers. In other words, some so-called nulls could really be positive or negative markers that remained unlabeled as such in those datasets even though the underlaying expression values indicated those patterns. This ambiguity about which designation is considered ‘true’ led to their removal from the classification assessment.

#### Creation of the Marker Standardization Workflow

The primary approach to standardizing marker names was to directly query the Protein Ontology. For these queries, the specified species was used to return both species-specific matches and matches that were unlinked to any species. When querying PRO, the inputted name was compared to the listed PRO term, PRO name, exact synonym, broad synonym, related synonym, and narrow synonym. Based on the quality of matches, ‘primary’ matches were considered inputs that connected to PRO terms, PRO names, and exact synonyms. ‘Secondary’ matches were the inputs that matched to broad, related, and narrow synonyms. If an input matched to both a primary and secondary term per species (with ‘no species’ considered a species type), only the primary matched term was returned. If an input only matched to the secondary term per species, then that term was outputted.

Due to the general lack of standardization of marker names used in the cytometry field, direct queries to PRO sometimes fail. As a result, other resources were added to the workflow to encompass more marker synonyms. It was observed that in many cases protein complexes (i.e. MHCII or TCRab) were not listed as PRO terms but rather GO terms. Because querying the GO directly could result in inaccurate GO term matches for the other protein markers, a list was created with all the GO complex terms used in the CL. This list was used in the workflow to automatically revert these protein complexes to GO terms. Additionally, a list of cluster of differentiation (CD) synonyms was used to automatically connect to additional marker synonyms[1]. If marker names were linked through this list, the algorithm would then query those names in PRO. Wikidata, which encompasses protein information from UniProt, InterPro, and the Protein Data Bank, was used as another reference[2–6]. SPARQL was used for these queries and if any synonyms were found then PRO would once again be queried.

When user-inputted marker names were not able to be automatically matched to a PRO term, the cause was often that the names were not specific enough. Automatically reverting to standardized marker names in this case may be making incorrect assumptions about the user's intentions. Instead, a list of suggestions was created by manually examining all the inputted marker names from the developmental dataset that failed. Some suggestions were taken from the list of CD synonyms, created by the Human Cell Differentiation Molecules organization, which was also used for the synonyms search[1]. This list was modified to include the broad CD identifiers that did not match and the more specific forms as suggestions. For example, an input of 'CD11' was too broad, so this list will suggest to the user they indicate if it is 'CD11a', 'CD11b', 'CD11c', or 'CD11d'. Other non-CD marker suggestions were added based on the development markers and how they were specified in PRO.

In addition to marker inputs not being specific enough, marker inputs often were in a format that made it impossible to provide an accurate match. For example, users frequently combined multiple markers together in their panel or referred to multiple markers as a non-protein term (i.e. 'lineage'). Additionally, user inputs sometimes included punctuation markers used in disparate ways, meaning they could not be removed or split universally (i.e. 'CD16a/b' and 'CD194/CCR4'). In these cases, the solution was to include suggestions to rewrite inputs when certain characters are used (commas, brackets, parentheses, and slashes). Lists of non-protein words that were commonly used were also created to tag inputs that either were not marker names at all (i.e. 'dead' or 'lineage') or still contained the metal or fluorochrome used in the experiment. The general 'non-protein' word list was created based off names found in the developmental set, with additional words added based off observed patterns (i.e. cytometry instrumentation names were added since it was observed they are sometimes connected to protein names). The 'fluorochrome' list was created based off fluorochromes listed in the 'reported_name' column of the downloaded 'reagent' file from ImmPort. Similarly, the 'metal tag' list was created based off metal ions listed in the 'reported_name' column of the same file. All curated suggestion lists are available on GitHub.

Since the commonly used markers CD3e, CD8a, and TCR are part of larger complexes, the workflow was created to include automatic matching to those complexes. Specifically, an input of CD3e matches to CD3e, TCR, TCRab, and TCRgd, while TCR matches to TCRab and TCRgd. CD8a matches to CD8a and CD8ab. If the more specific marker is also included in the query (TCRab, TCRgd, or CD8ab), this automatic matching will not occur.

The order of the workflow was determined based on maximizing the number of PRO or GO matches in the least amount of time. The final step of the workflow allows the user to incorporate any suggestions into renaming their unmatched markers.

### Figures

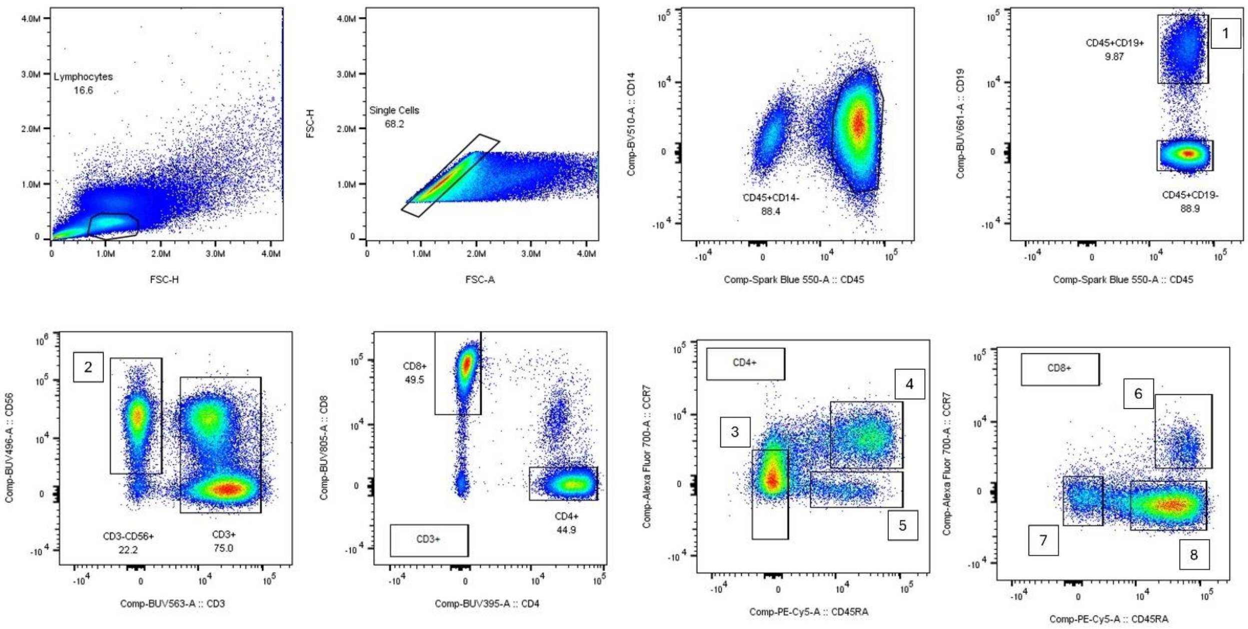


Figure S1. Manual gating scheme for the spectral flow cytometry dataset that was used for benchmarking the tool. The numbers indicate the populations and correspond to the following cell type names: 1) B cells, 2) Natural killer cells, 3) Effector memory $CD4^+$ T cells, 4) Naïve $CD4^+$ T cells, 5) TEMRA $CD4^+$ T cells, 6) Naïve $CD8^+$ T cells, 7) Effector memory $CD8^+$ T cells, and 8) TEMRA $CD8^+$ T cells.

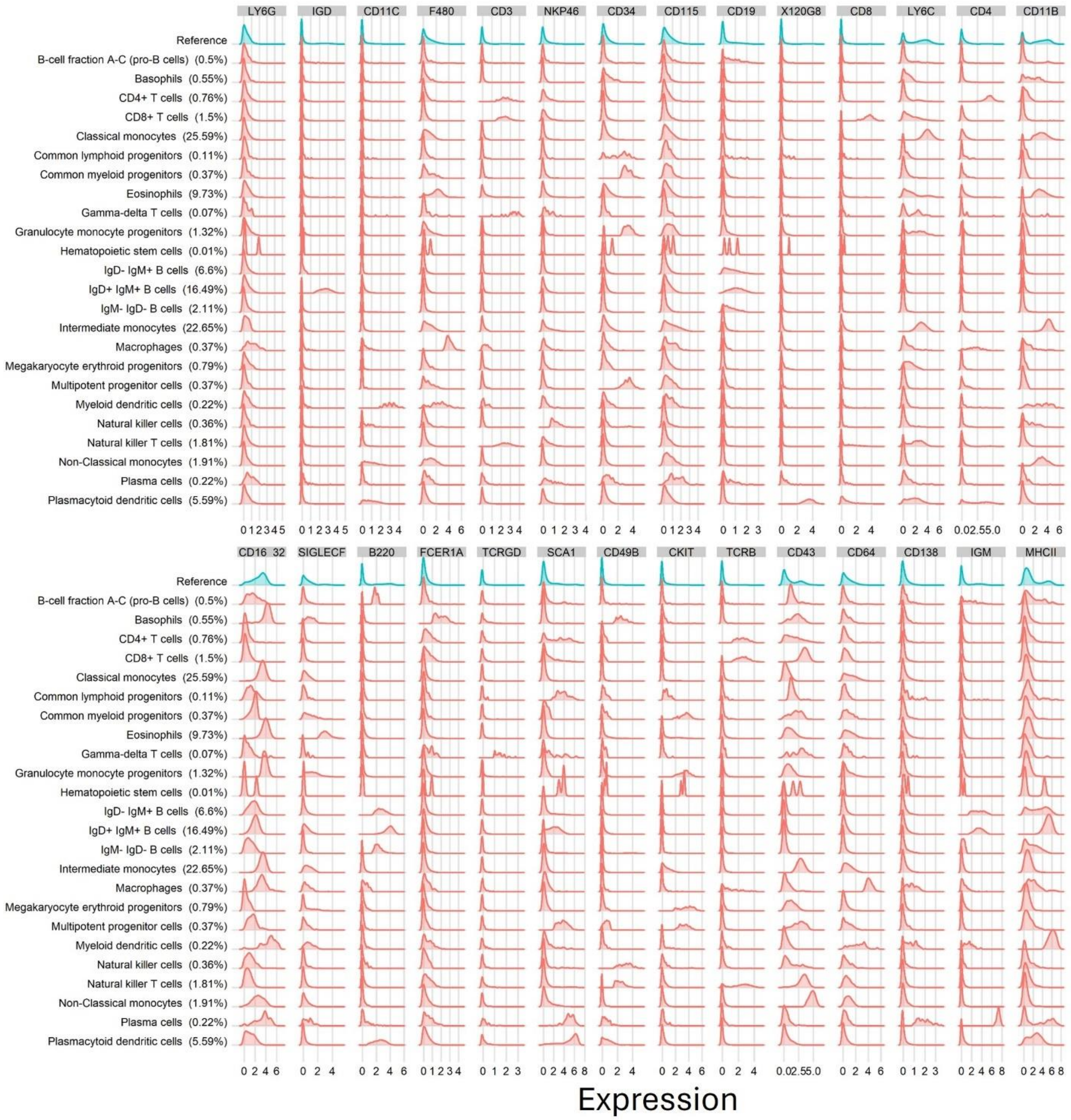


Figure S2. Histogram plots showing the expression distribution (arcsinh transformed) per cluster from the Samusik dataset[7]. Only markers used in the manual gating scheme are included. The percentage of live cells per cluster are shown in parentheses.

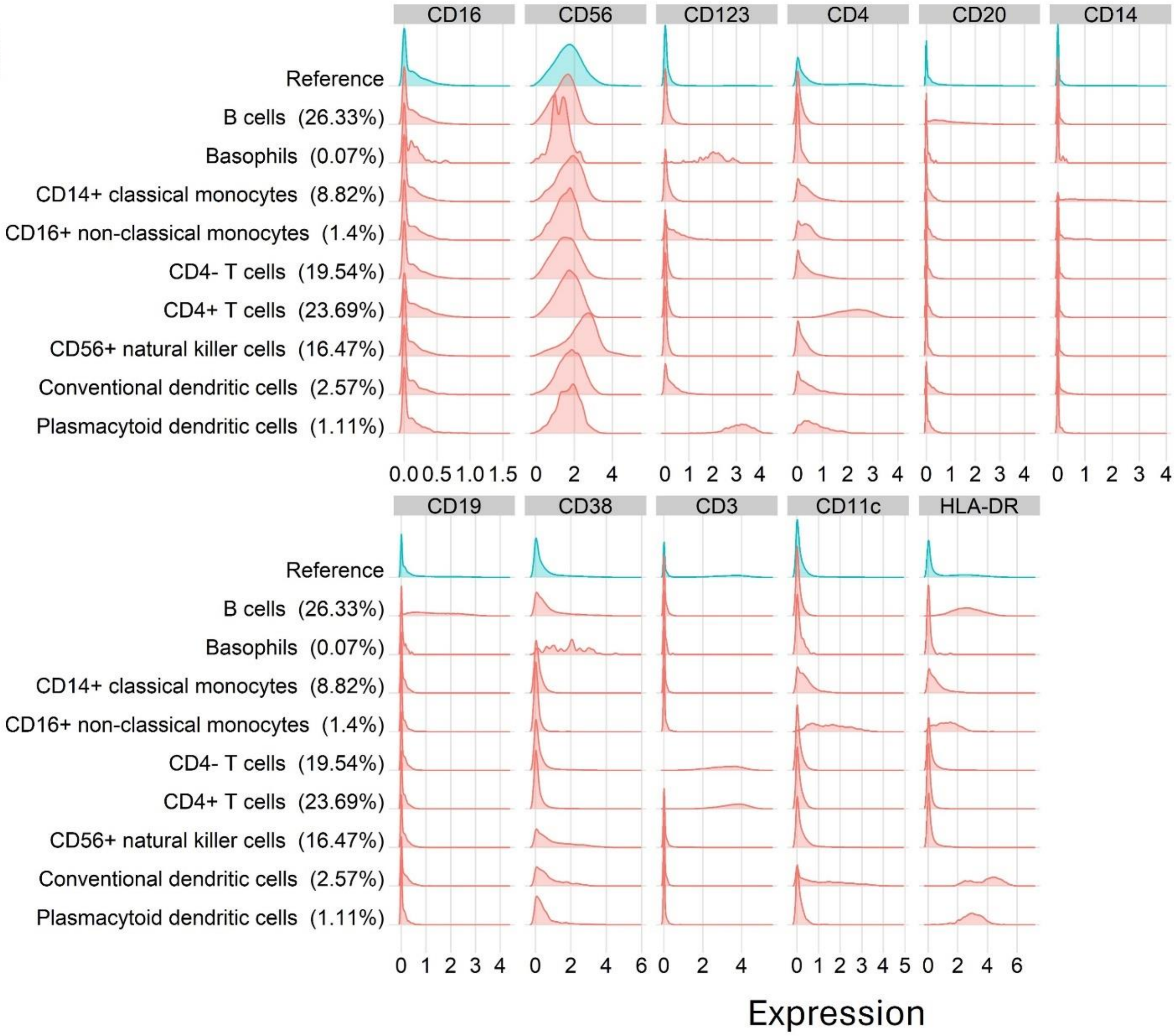


Figure S3. Histogram plots showing the expression distribution (arcsinh transformed) per cluster from the Kimmey dataset[8]. Only markers used in the cell type marker definitions are included. The percentage of live cells per cluster are shown in parentheses.

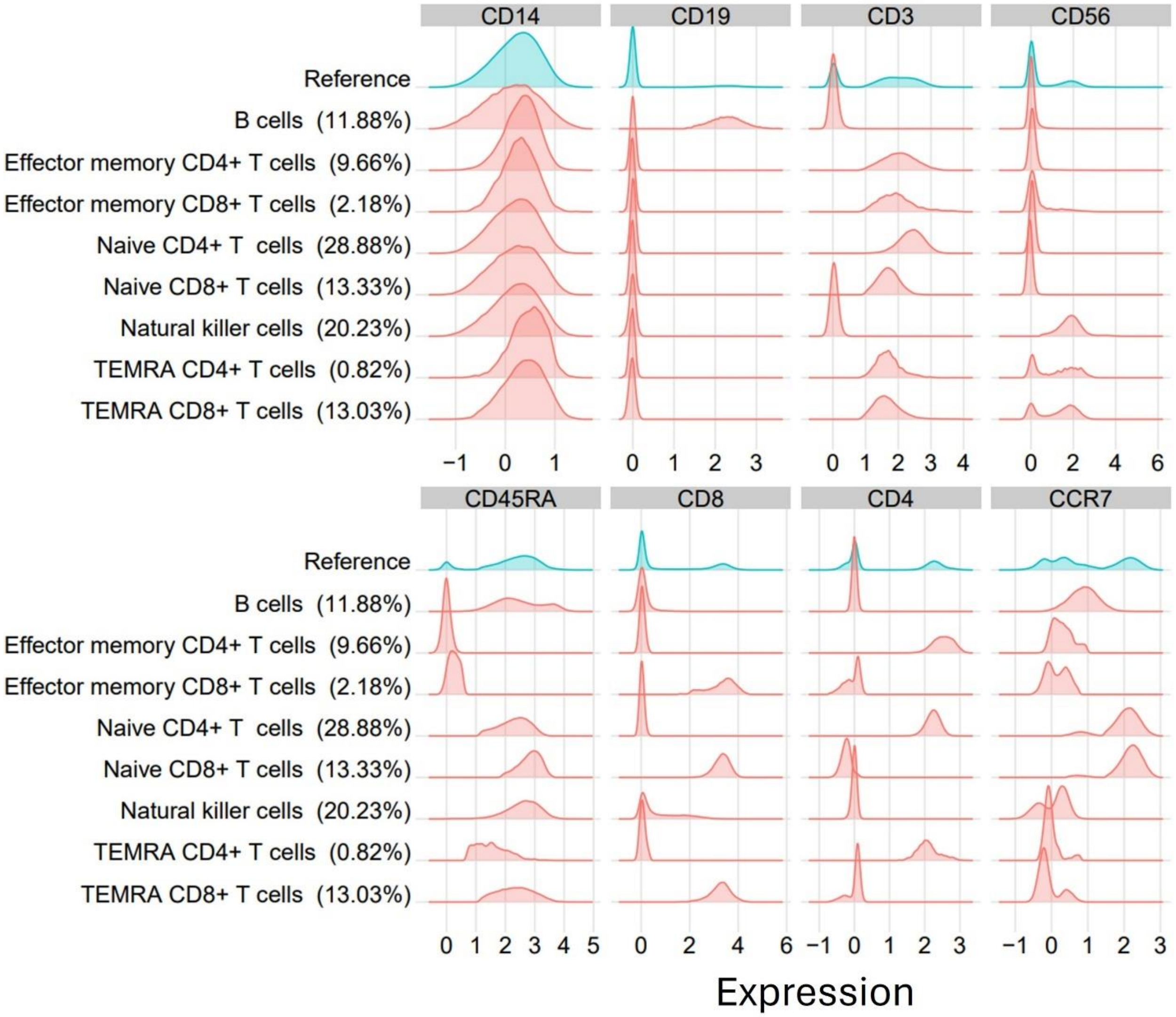


Figure S4. Histogram plots showing the expression distribution (arcsinh transformed) per cluster from the spectral dataset. Only markers used in the manual gating scheme are included. The percentage of live cells per cluster are shown in parentheses.

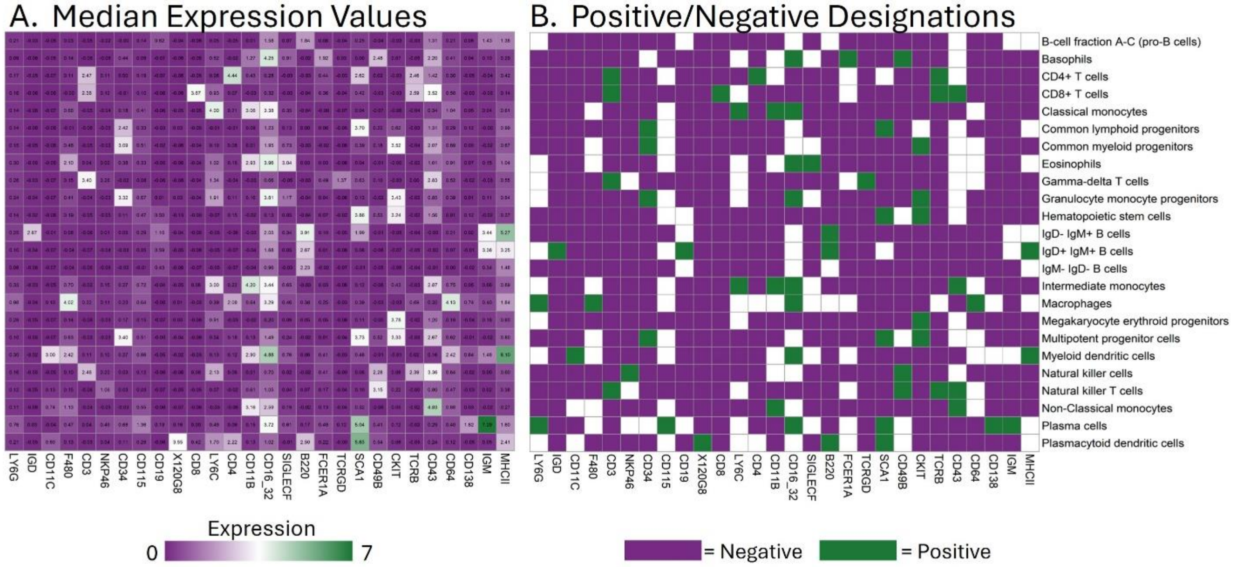


Figure S5. Heatmaps depicting the markers and cell type names that were delineated via manual gating analysis from Samusik et al[7]. Specifically, A) shows median expression values and B) shows results of the tool's marker designation algorithm (Part 1).

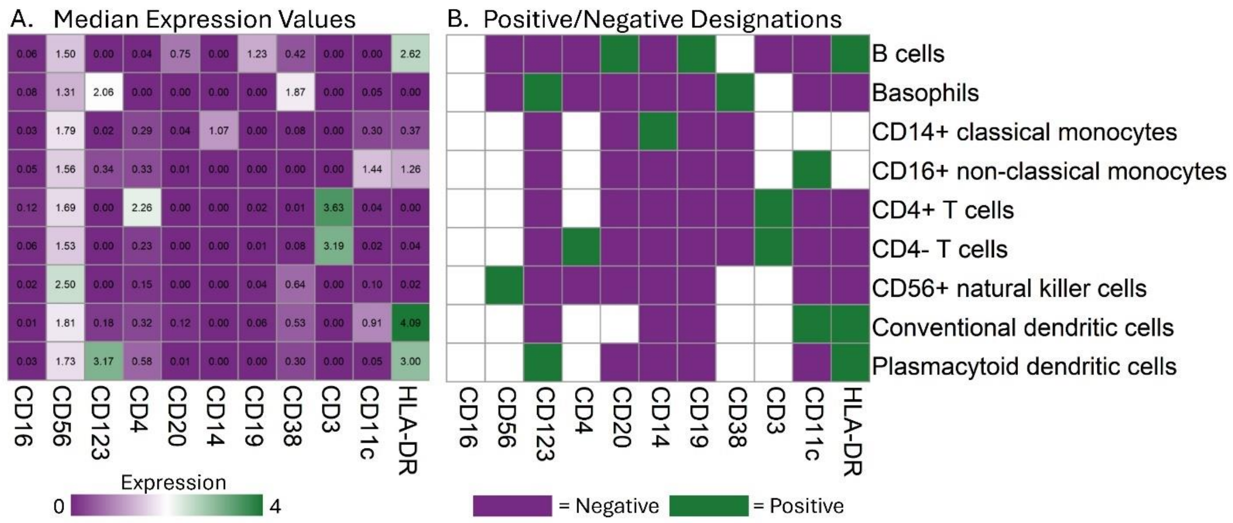


Figure S6. Heatmaps depicting the markers and cell type names that were delineated via clustering analysis from Kimmey et al[8]. Specifically, A) shows median expression values and B) shows results of the tool's marker designation algorithm (Part 1).

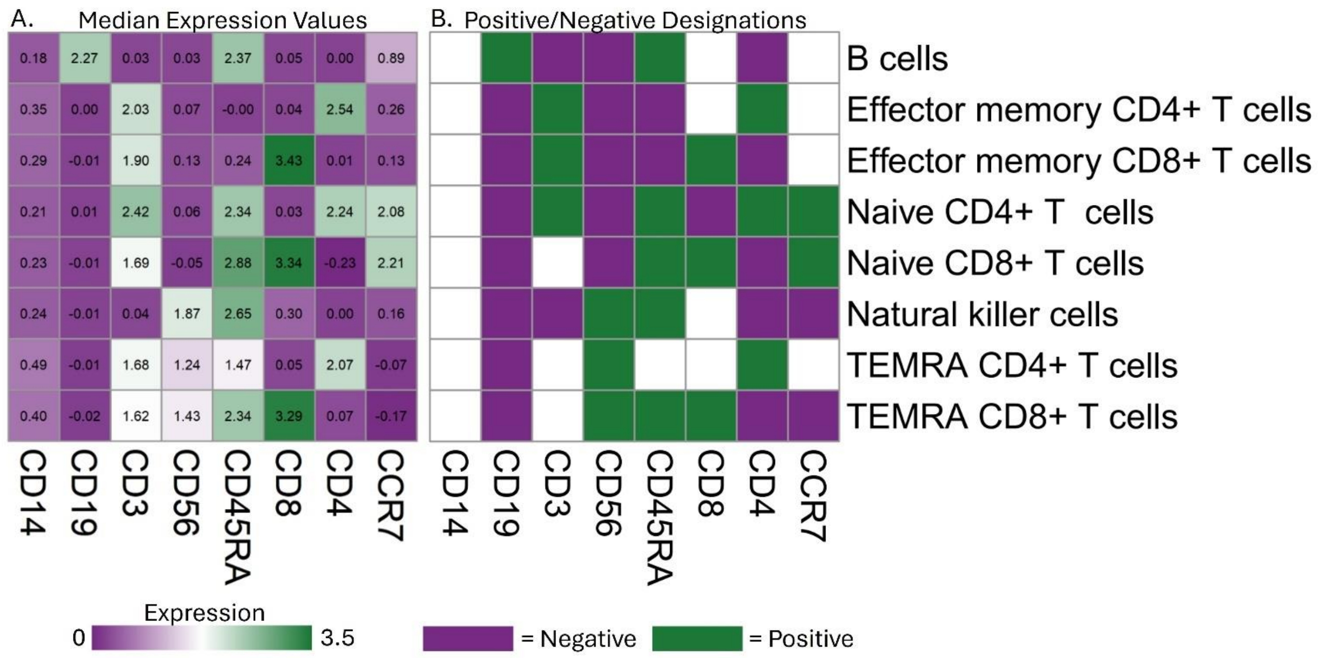


Figure S7. Heatmaps depicting the markers and cell type names that were delineated via manual gating analysis for the spectral flow cytometry data. Specifically, A) shows median expression values and B) shows results of the tool's marker designation algorithm (Part 1).

#### Tables

Table S1. Antibody panel used for the seven sample spectral cytometry dataset. Note that the fluorophores for CCR7 and HELIOS were switched for analysis of donors five through seven.

| Antigen | Fluorophore | Clone | Company |
|---|---|---|---|
| CD4 | BUV395 | M-T477 | BD Biosciences |
| CD56 | BUV496 | NCAM16.2 | BD Biosciences |
| CD3 | BUV563 | UCHT1 | BD Biosciences |
| CD19 | BUV661 | SJJJ25C1 | BD Biosciences |
| CD8 | BUV805 | RPA-T8 | BD Biosciences |
| CD103 | BV421 | Ber-ACT8 | Biolegend |
| FOXP3 | Pacific Blue | 259D | Biolegend |
| CD14 | BV510 | 63D3 | Biolegend |
| CD69 | BV605 | FN50 | Biolegend |
| CCR6 | BV650 | A019D5 | Biolegend |
| Perforin | BV711 | dG9 | Biolegend |
| CCR7 | AlexaFluor 488 (donors 5-7) | G043H7 | Biolegend |
| | AlexaFluor 700 (donors 1-4) | | |
| HELIOS | AlexaFluor 488 (donors 1-4) | 22F6 | Biolegend |
| | AlexaFluor 700 (donors 5-7) | | |
| CD45 | Spark Blue 550 | 2D1 | Biolegend |
| TIGIT | PerCP/eFluor710 | MBSA43 | eBiosciences |
| CD25 | PE | 2A3 | BD Biosciences |
| CD39 | PE/Dazzle594 | A1 | Biolegend |
| CD45RA | PE/Cy5 | HI100 | Biolegend |
| PD1 | PE/Cy7 | EH12.1 | BD Biosciences |
| CCR4 | APC | DT5D3 | Miltenyl Biotec |
| GZMB/GranzymeB | APC/Fire750 | QA16A02 | Biolegend |

Table S2. Summary of cell type names and marker definitions that were included in the bone marrow study published by Samusik et al. and used for benchmarking in this study[7]. All cell types that were listed by name in Supplementary Figure 5 from the original manuscript were included (N=24). The marker descriptions were taken from the manual gating strategy included in that supplemental figure. The equivalent CL name and term was manually determined for each original cell type based on the original name and the marker definition. Cell types that could not be matched to a CL term and were subsequently excluded from cell type matching tests are indicated.

| **Original cell type name** | **Original marker definitions** | **CL cell type name** | **CL term** |
|---|---|---|---|
| B-cell fraction A-C (pro-B cells) | $CD43^+$, $B220^+$, $120g8^-$ | No equivalent CL term - too broad | |
| Basophils | $FceR1a^+$, $NKp46^-$, $CD3^-$, $CD49b^+$, $B220^-$, $120g8^-$ | Basophil | CL_0000767 |
| $CD4^+$ T cells | $CD4^+$, $CD8^-$, $TCRb^+$, $TCRgd^-$, $CD3^+$, $CD49b^-$, $B220^-$, $120g8^-$ | CD4-positive, alpha-beta T cell | CL_0000624 |
| $CD8^+$ T cells | $CD8^+$, $CD4^-$, $TCRb^+$, $TCRgd^-$, $CD3^+$, $CD49b^-$, $B220^-$, $120g8^-$ | CD8-positive, alpha-beta T cell | CL_0000625 |
| Classical monocytes | $Ly6C^+$, $CD43^-$, $CD11b^+$, $CD3^-$, $CD49b^-$, $B220^-$, $120g8^-$ | Gr1-high classical monocyte | CL_0002395 |
| Common lymphoid progenitors | $CD43^+$, $B220^-$, $Sca1^+$, $cKit^+$, $CD115^-$, $CD11b^-$, $CD3^-$, $CD49b^-$, $B220^-$, $120g8^-$ | Kit-positive, Sca1-positive common lymphoid progenitor | CL_0001025 |
| Common myeloid progenitors | $CD34^+$, $CD16\_32^-$, $cKit^+$, $Sca1^-$, $CD115^-$, $CD11b^-$, $CD3^-$, $CD49b^-$, $B220^-$, $120g8^-$ | Kit-positive, CD34-positive common myeloid progenitor | CL_0001023 |
| Eosinophils | $SiglecF^+$, $CD11b^+$ | Eosinophil | CL_0000771 |
| Gamma-delta T cells | $TCRgd^+$, $TCRb^-$, $CD3^+$, $CD49b^-$, $B220^-$, $120g8^-$ | Gamma-delta T cell | CL_0000798 |
| Granulocyte monocyte progenitors | $CD16\_32^+$, $CD34^+$, $cKit^+$, $Sca1^-$, $CD115^-$, $CD11b^-$, $CD3^-$, $CD49b^-$, $B220^-$, $120g8^-$ | Kit-positive granulocyte monocyte progenitor | CL_0002002 |
| Hematopoietic stem cells | $CD34^-$, $cKit^+$, $Sca1^+$, $CD115^-$, $CD11b^-$, $CD3^-$, $CD49b^-$, $B220^-$, $120g8^-$ | Kit and Sca1-positive hematopoietic stem cell | CL_0001008 |
| $IgD^-$ $IgM^+$ B cells | $IgD^-$, $IgM^+$, $CD43^-$, $B220^+$, $120g8^-$ | Fraction E immature B cell | CL_0002054 |
| $IgD^+$ $IgM^+$ B cells | $IgD^+$, $IgM^+$, $CD43^-$, $B220^+$, $120g8^-$ | Fraction F mature B cell | CL_0002056 |
| $IgM^-$ $IgD^-$ B cells | $IgD^-$, $IgM^-$, $CD43^-$, $B220^+$, $120g8^-$ | Fraction D precursor B cell | CL_0002052 |

| | | | |
|---|---|---|---|
| Intermediate monocytes | Ly6C$^{+}$, CD43$^{+}$, CD11b$^{+}$, CD3$^{-}$, CD49b$^{-}$, B220$^{-}$, 120g8$^{-}$ | Gr1-positive, CD43-positive monocyte | CL_0002398 |
| Macrophages | F480$^{+}$, CD64$^{+}$, CD3$^{-}$, CD49b$^{-}$, B220$^{-}$, 120g8$^{-}$ | Bone marrow macrophage | CL_0002476 |
| Megakaryocyte erythroid progenitors | CD34$^{-}$, CD16_32$^{-}$, cKit$^{+}$, Sca1$^{+}$, CD115$^{-}$, CD11b$^{-}$, CD3$^{-}$, CD49b$^{-}$, B220$^{-}$, 120g8$^{-}$ | Kit-positive, CD34-negative megakaryocyte erythroid progenitor cell | CL_0002006 |
| Multipotent progenitor cells | CD34$^{+}$, DNA2$^{+}$, cKit$^{+}$, Sca1$^{+}$, CD115$^{-}$, CD11b$^{-}$, CD3$^{-}$, CD49b$^{-}$, B220$^{-}$, 120g8$^{-}$ | Slamf1-positive multipotent progenitor cell | CL_0002036 |
| | | Slamf1-negative multipotent progenitor cell | CL_0002035 |
| Myeloid dendritic cells | MHCII$^{+}$, CD11c$^{+}$, CD3$^{-}$, CD49b$^{-}$, B220$^{-}$, 120g8$^{-}$ | Mature CD16-positive myeloid dendritic cell | CL_0002534 |
| Natural killer cells | FceR1a$^{-}$, NKp46$^{+}$, CD3$^{-}$, CD49b$^{+}$, B220$^{-}$, 120g8$^{-}$ | Natural killer cell | CL_0000623 |
| Natural killer T cells | CD3$^{+}$, CD49b$^{+}$, B220$^{-}$, 120g8$^{-}$ | Mature NK T cell | CL_0000814 |
| | | Immature NK T cell stage IV | CL_0002042 |
| Non-Classical monocytes | Ly6C$^{-}$, CD43$^{+}$, CD11b$^{+}$, CD3$^{-}$, CD49b$^{-}$, B220$^{-}$, 120g8$^{-}$ | Gr1-low non-classical monocyte | CL_0002058 |
| Plasma cells | IgM$^{+}$, CD138$^{+}$ | Plasma cell | CL_0000786 |
| Plasmacytoid dendritic cells | 120g8$^{+}$, B220$^{+}$ | CD8_alpha-negative plasmacytoid dendritic cell | CL_0002455 |

Table S3. Summary of cell type names and marker definitions that were included in the PBMC study published by Kimmey et al. and used for benchmarking in this study[8]. All groups that were listed in Supplementary Table 3 from the original manuscript and were not used for *in silico* mononuclear cell purification (N=9) were included. Both the cell type names and marker descriptions were taken directly from that supplemental table. The equivalent CL name and term was manually determined for each original cell type based on the original name and the marker definition. Cell types that could not be matched to a CL term and were subsequently excluded from cell type matching tests are indicated. Markers that were designated as 'het' were excluded from analysis.

| **Original cell type name** | **Original marker definitions** | **CL cell type name** | **CL term** |
|---|---|---|---|
| B cells | $CD20^+$, $CD19^+$, $HLADR^+$ | No equivalent CL term - too broad | |
| Basophils | $CD38^+$, $CD123^+$, $CD19^-$, $HLADR^-$ | Basophil | CL_0000767 |
| $CD14^+$ classical monocytes | $CD14^+$, $CD16^{neg}$ | CD14-positive, CD16-negative classical monocyte | CL_0002057 |
| $CD16^+$ non-classical monocytes | $CD14^{low}$, $CD16^+$ | CD14-low, CD16-positive monocyte | CL_0002396 |
| $CD4^+$ T cells | $CD3^+$, $CD4^+$, $CD19^-$, $HLADR^-$ | CD4-positive, alpha-beta T cell | CL_0000624 |
| CD4- T cells | $CD3^+$, $CD4^-$, $CD19^-$, $HLADR^-$ | No equivalent CL term - too broad | |
| $CD56^+$ natural killer cells | $CD56^{++}$, $CD38^{het}$, $CD16^{het}$, $CD14^-$, $CD19^-$, $CD20^-$ | CD16-negative, CD56-bright natural killer cell, human | CL_0000938 |
| Conventional dendritic cells | $CD11c^+$, $HLADR^+$, CD14- | Immature conventional dendritic cell | CL_0000840 |
| | | Mature conventional dendritic cell | CL_0000841 |
| Plasmacytoid dendritic cells | $CD123^+$, $HLADR^+$, $CD14^-$ | Immature CD11c-negative plasmacytoid dendritic cell | CL_0000994 |

Table S4. Summary of cell type names and marker definitions that were included in the unpublished spectral cytometry PBMC study and used for benchmarking in this study. All cell types (N=8) that were numbered in the manual gating scheme were included. Marker descriptions were taken from the manual gating strategy. The equivalent CL name and term was manually determined for each original cell type based on the original name and the marker definition. Cell types that could not be matched to a CL term and were subsequently excluded from cell type matching tests are indicated.

| **Original cell type name** | **Original marker definitions** | **CL cell type name** | **CL term** |
|---|---|---|---|
| B cells | $CD14^-$, $CD19^+$ | B cell, CD19-positive | CL_0001201 |
| Effector memory $CD4^+$ T cells | $CD14^-$, $CD19^-$, $CD3^+$, $CD4^+$, $CD8^-$, $CD45RA^-$, $CCR7^-$ | Effector memory CD4-positive, alpha-beta T cell | CL_0000905 |
| Effector memory $CD8^+$ T cells | $CD14^-$, $CD19^-$, $CD3^+$, $CD4^-$, $CD8^+$, $CD45RA^-$, $CCR7^-$ | Effector memory CD8-positive, alpha-beta T cell | CL_0000913 |
| Naive $CD4^+$ T cells | $CD14^-$, $CD19^-$, $CD3^+$, $CD4^+$, $CD8^-$, $CD45RA^+$, $CCR7^+$ | Naive thymus-derived CD4-positive, alpha-beta T cell | CL_0000895 |
| Naive $CD8^+$ T cells | $CD14^-$, $CD19^-$, $CD3^+$, $CD4^-$, $CD8^+$, $CD45RA^+$, $CCR7^+$ | Naive thymus-derived CD8-positive, alpha-beta T cell | CL_0000900 |
| Natural killer cells | $CD14^-$, $CD19^-$, $CD3^-$, $CD56^+$ | CD16-negative, CD56-bright natural killer cell, human | CL_0000938 |
| | | CD16-positive, CD56-dim natural killer cell, human | CL_0000939 |
| TEMRA $CD4^+$ T cells | $CD14^-$, $CD19^-$, $CD3^+$, $CD4^+$, $CD8^-$, $CD45RA^+$, $CCR7^-$ | Effector memory CD4-positive, alpha-beta T cell, terminally differentiated | CL_0001087 |
| TEMRA $CD8^+$ T cells | $CD14^-$, $CD19^-$, $CD3^+$, $CD4^-$, $CD8^+$, $CD45RA^+$, $CCR7^-$ | Effector memory CD8-positive, alpha-beta T cell, terminally differentiated | CL_0001062 |

Table S5. Summary of cell type names and marker definitions that were included in OMIP 54 and used for benchmarking (Part 3) in this study[9]. All cell types that were numbered (N=9) in the original manuscript were included, while cell subtypes that represented activation states were excluded. Markers were defined through a combination of OMIP textual descriptions and from the manual gating strategy. The equivalent CL name and term was manually determined for each original cell type based on the original OMIP name and the marker definition. Cell types that could not be matched to a CL term and were subsequently excluded in testing are indicated.

| **Original cell type name** | **Original marker definitions** | **CL cell type name** | **CL term** |
|---|---|---|---|
| Dendritic cells ($CD11c^{high}$) | $CD11c^{high}$ | Conventional dendritic cell | CL_0000990 |
| $CD45^{-}$ cells | $CD45^{-}$ | No equivalent CL term - too broad | |
| CD8 T cells | $CD11b^{-}$, $CD3^{+}$, $TCRb^{+}$, $CD8^{+}$, $CD4^{-}$ | CD8-positive, alpha-beta T cell | CL_0000625 |
| CD4 T cells | $CD11b^{-}$, $CD3^{+}$, $TCRb^{+}$, $CD8^{-}$, $CD4^{+}$ | CD4-positive, alpha-beta T cell | CL_0000624 |
| Tregs | $CD11b^{-}$, $CD3^{+}$, $TCRb^{+}$, $CD8^{-}$, $CD4^{+}$, $FoxP3^{+}$ | CD4-positive, CD25-positive, alpha-beta regulatory T cell | CL_0000792 |
| Microglia | $CD11b^{+}$, $CD44^{-}$, $CD49d^{-}$ | Microglial cell | CL_0000129 |
| Granulocytes | $CD11b^{+}$, $Ly6G^{+}$ | Granulocyte | CL_0000094 |
| Monocytes | $CD11b^{+}$, $Ly6G^{-}$, $Ly6C^{high}$, $CCR2^{high}$, $CD49d^{+}$ | Gr1-high classical monocyte | CL_0002395 |
| Macrophages | $CD11b^{+}$, $Ly6G^{-}$ | Macrophage | CL_0000235 |

Table S6. Summary of cell type names and marker definitions that were included in OMIP 63 and used for benchmarking (Part 3) in this study[10]. All cell types that were listed in Table 3 (N=28) from the original manuscript were included. Markers were defined through a combination of OMIP textual descriptions and from the manual gating strategy. The equivalent CL name and term was manually determined for each original cell type based on the original OMIP name and the marker definition. Cell types that could not be matched to a CL term and were subsequently excluded in testing are indicated. Markers that were designated as ‘$^{+}$/$^{-}$’ and ‘low/$^{-}$’ were excluded from analysis.

| **Original cell type name** | **Original marker definitions** | **CL cell type name** | **CL term** |
|---|---|---|---|
| T cell | CD3$^{+}$, CD20$^{-}$ | No equivalent CL term - too broad | |
| CD8$^{+}$ T cell | CD3$^{+}$, CD20$^{-}$, CD8$^{+}$, CD4$^{-}$ | CD8-positive, alpha-beta T cell | CL_0000625 |
| Naïve CD8$^{+}$ T cell | CD3$^{+}$, CD20$^{-}$, CD8$^{+}$, CD4$^{-}$, CCR7$^{+}$, CD45RA$^{+}$ | Naive thymus-derived CD8-positive, alpha-beta T cell | CL_0000900 |
| Central memory CD8$^{+}$ T cell | CD3$^{+}$, CD20$^{-}$, CD8$^{+}$, CD4$^{-}$, CCR7$^{+}$, CD45RA$^{-}$ | Central memory CD8-positive, alpha-beta T cell | CL_0000907 |
| Effector memory CD8$^{+}$ T cell | CD3$^{+}$, CD20$^{-}$, CD8$^{+}$, CD4$^{-}$, CCR7$^{-}$, CD45RA$^{-}$ | Effector memory CD8-positive, alpha-beta T cell | CL_0000913 |
| Effector memory revertant CD8$^{+}$ T cell | CD3$^{+}$, CD20$^{-}$, CD8$^{+}$, CD4$^{-}$, CCR7$^{-}$, CD45RA$^{+}$ | Effector memory CD8-positive, alpha-beta T cell, terminally differentiated | CL_0001062 |
| CD4$^{+}$ T cell | CD3$^{+}$, CD20$^{-}$, CD8$^{-}$, CD4$^{+}$ | CD4-positive, alpha-beta T cell | CL_0000624 |
| CD4$^{+}$ regulatory T cell | CD3$^{+}$, CD20$^{-}$, CD8$^{-}$, CD4$^{+}$, CD25$^{high}$, CD127$^{low}$ | CD4-positive, CD25-positive, alpha-beta regulatory T cell | CL_0000792 |
| Naïve CD4$^{+}$ T cell | CD3$^{+}$, CD20$^{-}$, CD8$^{-}$, CD4$^{+}$, CCR7$^{+}$, CD45RA$^{+}$ | Naive thymus-derived CD4-positive, alpha-beta T cell | CL_0000895 |
| Central memory CD4$^{+}$ T cell | CD3$^{+}$, CD20$^{-}$, CD8$^{-}$, CD4$^{+}$, CCR7$^{+}$, CD45RA$^{-}$ | Central memory CD4-positive, alpha-beta T cell | CL_0000904 |
| Effector memory CD4$^{+}$ T cell | CD3$^{+}$, CD20$^{-}$, CD8$^{-}$, CD4$^{+}$, CCR7$^{-}$, CD45RA$^{-}$ | Effector memory CD4-positive, alpha-beta T cell | CL_0000905 |
| T follicular helper cell | CD3$^{+}$, CD20$^{-}$, CD8$^{-}$, CD4$^{+}$, CD25$^{low/-}$, CD127$^{+/-}$, CXCR5$^{+}$, CD45RA$^{-}$ | T follicular helper cell | CL_0002038 |
| T helper 1 cell | CD3$^{+}$, CD20$^{-}$, CD8$^{-}$, CD4$^{+}$, CD25$^{low/-}$, | T-helper 1 cell | CL_0000545 |

| | | | |
|---|---|---|---|
| | CD127$^{+/-}$, CXCR5$^{-}$, CD45RA$^{-}$, CXCR3$^{+}$, CCR6$^{-}$ | | |
| T helper 17 cell | CD3$^{+}$, CD20$^{-}$, CD8$^{-}$, CD4$^{+}$, CD25$^{low/-}$, CD127$^{+/-}$, CXCR5$^{-}$, CD45RA$^{-}$, CXCR3$^{-}$, CCR6$^{+}$ | T-helper 17 cell | CL_0000899 |
| Other T helper cells | CD3$^{+}$, CD20$^{-}$, CD8$^{-}$, CD4$^{+}$, CD25$^{low/-}$, CD127$^{+/-}$, CXCR5$^{-}$, CD45RA$^{-}$, CXCR3$^{-}$, CCR6$^{-}$ | No equivalent CL term - too broad | |
| B cell | CD20$^{+}$, CD3$^{-}$ | No equivalent CL term - too broad | |
| Transitional B cell | CD20$^{+}$, CD3$^{-}$, CD10$^{+}$, CD27$^{-}$ | Transitional stage B cell | CL_0000818 |
| Naïve B cell | CD20$^{+}$, CD3$^{-}$, CD10$^{-}$, CD27$^{-}$ | Naive B cell | CL_0000788 |
| Memory B cell | CD20$^{+}$, CD3$^{-}$, CD10$^{-}$, CD27$^{+}$ | Memory B cell | CL_0000787 |
| Atypical or aged memory B cell | CD20$^{+}$, CD3$^{-}$, CD19$^{high}$, CD21$^{low}$ | No equivalent CL term - not in CL | |
| IgM$^{+}$ memory B cell | CD10$^{-}$, CD27$^{+}$, IgM$^{+}$, IgG$^{-}$, IgA$^{-}$ | IgM memory B cell | CL_0000971 |
| IgG$^{+}$ switched memory B cell | CD10$^{-}$, CD27$^{+}$, IgM$^{-}$, IgG$^{+}$, IgA$^{-}$ | IgG memory B cell | CL_0000979 |
| IgA$^{+}$ switched memory B cell | CD10$^{-}$, CD27$^{+}$, IgM$^{-}$, IgG$^{-}$, IgA$^{+}$ | IgA memory B cell | CL_0000973 |
| Natural killer cell | CD3$^{-}$, CD56$^{+}$ | Natural killer cell | CL_0000623 |
| Unconventional T cells (CD3$^{+}$CD20$^{-}$) | CD20$^{-}$, CD3$^{+}$ | No equivalent CL term - too broad | |
| Natural killer T cell | CD20$^{-}$, CD3$^{+}$, TCR Vα24JαQ$^{+}$ | Mature NK T cell | CL_0000814 |
| Mucosal-associated invariant T cell | CD20$^{-}$, CD3$^{+}$, CD161$^{+}$, TCR Vα7.2$^{+}$ | Mucosal invariant T cell | CL_0000940 |
| Gamma-delta T cell | CD20$^{-}$, CD3$^{+}$, TCR Vab$^{-}$, TCR Vgd$^{+}$ | Gamma-delta T cell | CL_0000798 |

Table S7. Summary of cell type names and marker definitions that were included in OMIP 78 and used for benchmarking (Part 3) in this study[11]. All cell types that were numbered (N=25) in the original manuscript were included. Markers were defined through a combination of OMIP textual descriptions and from the manual gating strategy. The equivalent CL name and term was manually determined for each original cell type based on the original OMIP name and the marker definition. Cell types that could not be matched to a CL term and were subsequently excluded in testing are indicated.

| **Original cell type name** | **Original marker definitions** | **CL cell type name** | **CL term** |
|---|---|---|---|
| Naïve $CD4^+$ T cell | $CD45^+$, $CD19^-$, $CD14^-$, $CD56^-$, $CD16^-$, $CD133^-$, $CD3^+$, $CD4^+$, $CD8^-$, $CD27^+$, $CD45RO^-$ | Naive thymus-derived CD4-positive, alpha-beta T cell | CL_0000895 |
| Effector/effector memory $CD4^+$ T cell | $CD45^+$, $CD19^-$, $CD14^-$, $CD56^-$, $CD16^-$, $CD133^-$, $CD3^+$, $CD4^+$, $CD8^-$, $CD27^-$ | Effector CD4-positive, alpha-beta T cell | CL_0001044 |
| | | Effector memory CD4-positive, alpha-beta T cell | CL_0000905 |
| Central memory $CD4^+$ T cell | $CD45^+$, $CD19^-$, $CD14^-$, $CD56^-$, $CD16^-$, $CD133^-$, $CD3^+$, $CD4^+$, $CD8^-$, $CD27^+$, $CD45RO^+$ | Central memory CD4-positive, alpha-beta T cell | CL_0000904 |
| Naïve $CD8^+$ T cell | $CD45^+$, $CD19^-$, $CD14^-$, $CD56^-$, $CD16^-$, $CD133^-$, $CD3^+$, $CD4^-$, $CD8^+$, $CD27^+$, $CD45RO^-$ | Naive thymus-derived CD8-positive, alpha-beta T cell | CL_0000900 |
| Effector/effector memory $CD8^+$ T cell | $CD45^+$, $CD19^-$, $CD14^-$, $CD56^-$, $CD16^-$, $CD133^-$, $CD3^+$, $CD4^-$, $CD8^+$, $CD27^-$ | Effector CD8-positive, alpha-beta T cell | CL_0001050 |
| | | Effector memory CD8-positive, alpha-beta T cell | CL_0000913 |
| Central memory $CD8^+$ T cell | $CD45^+$, $CD19^-$, $CD14^-$, $CD56^-$, $CD16^-$, $CD133^-$, $CD3^+$, $CD4^-$, $CD8^+$, $CD27^+$, $CD45RO^+$ | Central memory CD8-positive, alpha-beta T cell | CL_0000907 |
| Natural killer T cell | $CD45^+$, $CD19^-$, $CD14^-$, $CD56^+$, $CD3^+$ | Mature NK T cell | CL_0000814 |
| $CD16^-$ $CD56^+$ natural killer cell | $CD45^+$, $CD19^-$, $CD14^-$, $CD3^-$, $CD16^-$, $CD56^+$ | CD16-negative, CD56-bright natural killer cell, human | CL_0000938 |
| $CD16^+$ $CD56^{low}$ natural killer cell | $CD45^+$, $CD19^-$, $CD14^-$, $CD3^-$, $CD16^+$, $CD56^{low}$ | CD16-positive, CD56-dim natural killer cell, human | CL_0000939 |

| | | | |
|---|---|---|---|
| $CD16^{low}$ $CD56^{+}$ natural killer cell | $CD45^{+}$, $CD19^{-}$, $CD14^{-}$, $CD3^{-}$, $CD16^{low}$, $CD56^{+}$ | No equivalent CL term - not in CL | |
| $CD16^{+}$ $CD56^{-}$ natural killer cell | $CD45^{+}$, $CD19^{-}$, $CD14^{-}$, $CD3^{-}$, $CD16^{+}$, $CD56^{-}$ | No equivalent CL term - not in CL | |
| Hematopoietic progenitor cell | $CD45^{+}$, $CD19^{-}$, $CD14^{-}$, $CD56^{-}$, $CD3^{-}$, $CD11b^{-}$, HLA-$DR^{-}$, $CD34^{+}$ | Hematopoietic multipotent progenitor cell | CL_0000837 |
| Naive B cell | $CD45^{+}$, $CD14^{-}$, $CD56^{-}$, $CD3^{-}$, $CD19^{+}$, $CD27^{-}$, $IgD^{+}$ | Naive B cell | CL_0000788 |
| Unswitched memory B cell | $CD45^{+}$, $CD14^{-}$, $CD56^{-}$, $CD3^{-}$, $CD19^{+}$, $CD27^{+}$, $IgD^{+}$ | Unswitched memory B cell | CL_0000970 |
| Memory B cell | $CD45^{+}$, $CD14^{-}$, $CD56^{-}$, $CD3^{-}$, $CD19^{+}$, $CD27^{+}$, $IgD^{-}$ | IgD-negative memory B cell | CL_0001053 |
| $IgG^{+}$ memory B cell | $CD45^{+}$, $CD14^{-}$, $CD56^{-}$, $CD3^{-}$, $CD19^{+}$, $CD27^{+}$, $IgD^{-}$, $IgM^{-}$, $IgG^{+}$ | IgG memory B cell | CL_0000979 |
| Plasma cell | $CD45^{+}$, $CD14^{-}$, $CD56^{-}$, $CD3^{-}$, $CD19^{+}$, $CD27^{+}$, $IgD^{-}$, $IgM^{-}$, $CD38^{+}$, $CD138^{-}$ | Plasma cell | CL_0000786 |
| Plasmablast | $CD45^{+}$, $CD14^{-}$, $CD56^{-}$, $CD3^{-}$, $CD19^{+}$, $CD27^{+}$, $IgD^{-}$, $IgM^{-}$, $CD38^{+}$, $CD138^{+}$ | Plasmablast | CL_0000980 |
| Transitional stage B cell | $CD45^{+}$, $CD14^{-}$, $CD56^{-}$, $CD3^{-}$, $CD19^{+}$, $CD27^{-}$, $CD38^{+}$, $CD24^{+}$ | Transitional stage B cell | CL_0000818 |
| Plasmacytoid dendritic cell | $CD45^{+}$, $CD14^{-}$, $CD3^{-}$, $CD19^{-}$, $CD56^{-}$, $CD11b^{-}$, HLA-$DR^{+}$, $CD11c^{-}$, $CD123^{+}$, $CD303^{+}$ | Plasmacytoid dendritic cell, human | CL_0001058 |
| $CD141^{+}$ conventional dendritic cell | $CD45^{+}$, $CD14^{-}$, $CD3^{-}$, $CD19^{-}$, $CD56^{-}$, $CD11b^{-}$, HLA-$DR^{+}$, $CD11c^{+}$, $CD1c^{-}$, $CD141^{+}$ | CD141-positive myeloid dendritic cell | CL_0002394 |

| | | | |
|---|---|---|---|
| $CD1c^+$ conventional dendritic cell | $CD45^+$, $CD14^-$, $CD3^-$, $CD19^-$, $CD56^-$, $CD11b^-$, $HLA\text{-}DR^+$, $CD11c^+$, $CD1c^+$, $CD141^-$ | CD1c-positive myeloid dendritic cell | CL_0002399 |
| Classical monocyte | $CD45^+$, $CD3^-$, $CD19^-$, $CD56^-$, $CD8^-$, $CD34^-$, $HLA\text{-}DR^+$, $CD14^+$, $CD16^-$ | CD14-positive, CD16-negative classical monocyte | CL_0002057 |
| Intermediate monocyte | $CD45^+$, $CD3^-$, $CD19^-$, $CD56^-$, $CD8^-$, $CD34^-$, $HLA\text{-}DR^+$, $CD14^+$, $CD16^+$ | CD14-positive, CD16-low monocyte | CL_0001055 |
| Non-classical monocyte | $CD45^+$, $CD3^-$, $CD19^-$, $CD56^-$, $CD8^-$, $CD34^-$, $HLA\text{-}DR^+$, $CD14^-$, $CD16^+$ | Non-classical monocyte | CL_0000875 |

Table S8. Studies from FlowRepository and the ImmPort repository that were used to extract user-inputted marker names for the benchmarking of the marker standardization workflow (Part 2). The ImmPort or FlowRepository study accession number, the listed study name, and the species used in the study are shown.

| ImmPort (SDY) or FlowRepository (FR) Study Accession | Listed title | Species |
|---|---|---|
| SDY1530 | ZIKV ex vivo PBMCs study | Homo sapiens |
| SDY1733 | Single-Cell immune signature for detecting early-stage HCC and early assessing PD-1 immunotherapy efficacy | Homo sapiens |
| SDY1393 | Toll-like Receptors in Older Adults and Response to Vaccination - Year2 | Homo sapiens |
| SDY1396 | Systems Immune Profiling of Divergent Responses to Infection | Homo sapiens |
| SDY1397 | Systems Immune Profiling of Divergent Responses to Infection - Year2 | Homo sapiens |
| SDY1394 | Blood and tissue studies of Tick-borne diseases in adults. Blood and tissue studies of Tick-borne diseases in children. Year1 | Homo sapiens |
| SDY1395 | Blood and tissue studies of Tick-borne diseases in adults. Blood and tissue studies of Tick-borne diseases in children. Year2 | Homo sapiens |
| SDY1998 | Stereotactic Body Radiation Therapy in Treating Patients With Metastatic Kidney Cancer Undergoing Surgery | Homo sapiens |
| SDY1538 | Systems Biology to Identify Biomarkers of Neonatal Vaccine Immunogenicity | Homo sapiens |
| SDY2011 | A shift in lung macrophage composition is associated with COVID-19 severity and recovery | Homo sapiens |
| SDY2107 | Tissue adaptation and clonal segregation of human memory T cells in barrier sites | Homo sapiens |
| SDY2075 | Deficiency of PD-L1$^{+}$ Non-Immune Cells in Preterm Labor | Homo sapiens |
| SDY2159 | Cytotoxic T cells targeting spike glycoprotein are associated with hybrid immunity to SARS-CoV-2 | Homo sapiens |
| SDY2060 | Immune memory to COVID-19 vaccines | Homo sapiens |
| SDY2365 | Interactions between Siglec-8 and endogenous sialylated cis ligands restrain cell death induction in human eosinophils and mast cells | Homo sapiens |
| FR-FCM-Z6J8 | in vivo chemotherapy mediated cancer dormancy escape and its prevention | Mus musculus |
| FR-FCM-Z64U | MYC induced AML model | Mus musculus |

| | | |
|---|---|---|
| FR-FCM-Z63E | OMIP-0XX: 40-Color Spectral Flow Cytometry Delineates All Major Leukocyte Populations in Murine Lymphoid Tissues | Mus musculus |
| FR-FCM-Z4GN | Tissue resident lymphocyte landscape of murine lungs post recovery from pneumococcal pneumonia | Mus musculus |
| FR-FCM-Z4TE | Fetal liver autofluorescence | Mus musculus |
| FR-FCM-Z56F | Development and characterization of a mass cytometry panel for detecting the effect of acute doxorubicin exposure on murine cardiac non-myocytes | Mus musculus |
| FR-FCM-Z4LQ | Surface phenotypes of naïve and memory B cells in murine tissues | Mus musculus |
| FR-FCM-Z4Q2 | 27-color murine lung immunophenotyping (unmixed data) | Mus musculus |
| FR-FCM-Z4NB | A 33-color panel for phenotypic analysis of murine organ specific immune cells | Mus musculus |
| FR-FCM-Z3MY | Tumor infiltrating lymphocyes from Arid5a KO or WT murine tumors | Mus musculus |

Table S9. Contingency table depicting the binary classifier used to assess the results of the median difference equation (Part 1). The rows correspond to actual values and the columns correspond to classified/predicted values.

| | | **Predicted condition** | |
|---|---|---|---|
| | | **Predicted Positive** | **Predicted Negative** |
| **Actual condition** | **Positive** | Positive markers to positive markers | Positive markers to null markers and positive markers to negative markers |
| | **Negative** | Negative markers to null markers and negative markers to positive markers | Negative markers to negative markers |

Table S10. Studies from the ImmPort repository that were used to extract user-inputted marker names for the development of the marker standardization workflow (Part 2). The ImmPort study accession number, the ImmPort listed study name, and the species used in the study are shown.

| **ImmPort Study Accession** | **Listed title** | **Species** |
|---|---|---|
| SDY1468 | B-cell Immunity to Influenza (SLVP017) 2010 | Homo sapiens |
| SDY1471 | B-cell Immunity to Influenza (SLVP017) 2013 | Homo sapiens |
| SDY1466 | Monozygotic and Dizygotic Twin Pair T-Cell Responses to Influenza Vaccination SLVP018 2013 | Homo sapiens |
| SDY1469 | B-cell Immunity to Influenza (SLVP017) 2011 | Homo sapiens |
| SDY1464 | T cell responses to H1N1v and a longitudinal study of seasonal influenza vaccination SLVP015 2014 | Homo sapiens |
| SDY311 | T cell responses to H1N1v and a longitudinal study of seasonal influenza vaccination (TIV) SLVP015 2010 (See companion studies SDY315 2012 / SDY312 2009 / SDY314 2008 / SDY112 2011) | Homo sapiens |
| SDY112 | T cell responses to H1N1v and a longitudinal study of seasonal influenza vaccination (TIV) SLVP015 2011 (See companion studies SDY311 2010 / SDY312 2009 / SDY314 2008 / SDY315 2012) | Homo sapiens |
| SDY515 | Monozygotic and Dizygotic Twin Pair T-Cell Responses to Influenza Vaccination SLVP018 2010 | Homo sapiens |
| SDY519 | Monozygotic and Dizygotic Twin Pair T-Cell Responses to Influenza Vaccination SLVP018 2011 | Homo sapiens |
| SDY113 | Plasmablast response to inactivated and live attenuated influenza vaccines (TIV3/TIV3 ID/LAIV) SLVP021 2011 | Homo sapiens |
| SDY312 | T cell responses to H1N1v and a longitudinal study of seasonal influenza vaccination (TIV) SLVP015 2009 (See companion studies SDY315 2012 / SDY314 2008 / SDY311 2010 / SDY112 2011) | Homo sapiens |
| SDY314 | T cell responses to H1N1v and a longitudinal study of seasonal influenza vaccination (TIV) SLVP015 2008 (See companion studies SDY315 2012 / SDY312 2009 / SDY311 2010 / SDY112 2011) | Homo sapiens |
| SDY364 | Systems Biology Approach to Study Influenza Vaccine 2012-13 in Healthy Children (see companion studies SDY144, SDY368, SDY387, SDY522) | Homo sapiens |
| SDY368 | Systems Biology Approach to Study Influenza Vaccine 2013-14 in Healthy Children (see companion studies SDY364, SDY144, SDY387, SDY522) | Homo sapiens |
| SDY369 | Systems Biology Approach to Study Influenza Vaccine in Children with Autoimmunity (Juvenile | Homo sapiens |

| | | |
|---|---|---|
| | Dermatomyositis JDM) 2011/2012 Cohort (see companion studies SDY376, SDY372, SDY645) | |
| SDY144 | Systems Biology Approach to Study Influenza Vaccine 2011-12 in Healthy Children (see companion studies SDY364, SDY368, SDY387, SDY522) | Homo sapiens |
| SDY372 | Systems Biology Approach to Study Influenza Vaccine in Children with Autoimmunity (Juvenile Dermatomyositis JDM) 2012/2013 Cohort (see companion studies (SDY369, SDY376, SDY645) | Homo sapiens |
| SDY376 | Systems Biology Approach to Study Influenza Vaccine in Children with Autoimmunity (Juvenile Dermatomyositis JDM) 2013/2014 Cohort (see companion studies SDY369, SDY372, SDY645) | Homo sapiens |
| SDY387 | Systems Biology Approach to Study Influenza Vaccine 2010-11 in Healthy Children (see companion studies SDY144, SDY368, SDY387, SDY522) | Homo sapiens |
| SDY416 | Study to measure the immune response to the influenza vaccine in patients with chronic plaque psoriasis | Homo sapiens |
| SDY396 | Immune Responses to Seasonal LAIV 2011-2012 Influenza Vaccination in Humans (see companion study SDY224,SDY564) | Homo sapiens |
| SDY315 | T cell responses to H1N1v and a longitudinal study of seasonal influenza vaccination (TIV) SLVP015 2012 (See companion studies SDY311 2010 / SDY312 2009 / SDY314 2008 / SDY112 2011) | Homo sapiens |
| SDY461 | Monitoring of tissue-specific immune responses in man to naturally occurring pathogens using mass cytometric monitoring | Homo sapiens |
| SDY305 | Plasmablast response to inactivated and live attenuated influenza vaccines (TIV3/TIV3 ID) SLVP021 2012 | Homo sapiens |
| SDY472 | Plasmablast response to inactivated and live attenuated influenza vaccines (TIV3/TIV3 ID) in SLVP021 2013 | Homo sapiens |
| SDY475 | Characterization of the human maternal immune response 6 months or greater post delivery via cytometry Time-of-Flight(CyTOF). | Homo sapiens |
| SDY202 | Heterovariant cross-reactive B-cell responses induced by the 2009 pandemic influenza virus A subtype H1N1 vaccine SLVP021 | Homo sapiens |
| SDY984 | Zoster vaccine in young and elderly | Homo sapiens |

| | | |
|---|---|---|
| SDY56 | Systems Biology of 2010 trivalent Influenza vaccine (TIV) in young and elderly (see companion study SDY61 2007, SDY270 2009, SDY119 2011) | Homo sapiens |
| SDY478 | T cell responses to H1N1v and a longitudinal study of seasonal influenza vaccination SLVP015 2013 | Homo sapiens |
| SDY506 | CD107 CTL assay | Homo sapiens |
| SDY58 | Defining signatures for immune responsiveness by functional systems immunology | Homo sapiens |
| SDY517 | Natural Killer cells in resistance to infection with West Nile virus | Homo sapiens |
| SDY404 | Immunologic and genomic signatures of influenza vaccine response - 2011 (see companion studies SDY63, SDY400, SDY520) | Homo sapiens |
| SDY67 | Bioinformatics Approach to 2010-2011 TIV Influenza A/H1N1 Vaccine Immune Profiling | Homo sapiens |
| SDY420 | Immunobiology of Aging | Homo sapiens |
| SDY565 | IL2-RAPA ITN018AI: Proleukin and Rapamune in Type 1 Diabetes Mellitus | Homo sapiens |
| SDY622 | Humoral responses to Influenza vaccination in aged populations - Year 2 2012 (See companion studies SDY272 2011, SDY648 2013, SDY739 2014, SDY819 2015) | Homo sapiens |
| SDY645 | Systems Biology Approach to Study Influenza Vaccine in Children with Autoimmunity (Juvenile Dermatomyositis JDM) 2014/2015 Cohort (see companion studies SDY369, SDY376, SDY372) | Homo sapiens |
| SDY80 | Cellular and molecular characterization of the immune response in healthy NIH employees at baseline and after immunization with the H1N1 or seasonal influenza vaccines | Homo sapiens |
| SDY564 | Immune Responses to Seasonal TIV 2012-2013 Influenza Vaccination in Humans (see companion study SDY396,SDY224) | Homo sapiens |
| SDY89 | Systems Biology Analysis of the response to Licensed Hepatitis B Vaccine (Engerix-B) (see companion study SDY690) | Homo sapiens |
| SDY702 | Human T Cell Profile | Homo sapiens |
| SDY1097 | Early-life compartmentalization of human T cells | Homo sapiens |
| SDY1041 | CMV CD8 T Cells | Homo sapiens |
| SDY736 | Human Aging and CMV | Homo sapiens |
| SDY720 | Age-related alterations in innate immune responses (See companion study SDY736) | Homo sapiens |
| SDY788 | Immune Profiles to Predict Response to Desensitization Therapy in Highly HLA-Sensitized Kidney Transplant Candidates | Homo sapiens |

| | | |
|---|---|---|
| SDY648 | Humoral responses to Influenza vaccination in aged populations - Year 3 2013 (See companion studies SDY272 2011, SDY622 2012, SDY739 2014, SDY819 2015) | Homo sapiens |
| SDY887 | Defective signaling in aging, influenza vaccination 2007 SLVP015 | Homo sapiens |
| SDY739 | Humoral responses to Influenza vaccination in aged populations - Year 4 2014 (See companion studies SDY272 2011, SDY622 2012, SDY648 2013, SDY819 2015) | Homo sapiens |
| SDY820 | Human Immune Signature of Mycobacterium Tuberculosis infection. (See SDY1324 for sorted cell gene expression data) | Homo sapiens |
| SDY1324 | Transcriptomic Analysis of $CD4^+$ T Cells Reveals Novel Immune Signatures of Latent Tuberculosis. (See SDY820 for corresponding PBMC results.) | Homo sapiens |
| SDY888 | Human Immune Signature of Dengue virus infection- Gene Expression of CD4 subsets | Homo sapiens |
| SDY997 | AMP Lupus Network Project: Molecular Characterization of Lupus Nephritis and Correlation with Response to Therapy | Homo sapiens |
| SDY998 | AMP Rheumatoid Arthritis Phase 1 | Homo sapiens |
| SDY194 | Innate Immune Pathways in Elderly and Immunosuppressed Populations 2011 | Homo sapiens |
| SDY819 | Humoral responses to Influenza vaccination in aged populations - Year 5 2015 (See companion studies SDY272 2011, SDY622 2012, SDY648 2013, SDY739 2014) | Homo sapiens |
| SDY901 | Activation of the PD-1 Pathway Contributes to Immune Escape in EGFR-Driven Lung Tumors | Homo sapiens |
| SDY824 | Anti-TNF Agents in RA (ARA06) | Homo sapiens |
| SDY1100 | Project 3 - Optimization experiments DCs DV2 and DV4 - CyTOF | Homo sapiens |
| SDY1486 | Development of a Comprehensive Antibody Staining Database using a Standardized Analytics Pipeline | Homo sapiens |
| SDY1594 | Modified CyTOF Helios Injector Can Improve Data Quality | Homo sapiens |
| SDY1115 | Comparative analysis of neonatal and adult immune responses after in vitro rhinovirus stimulation | Homo sapiens |
| SDY1157 | Immune response throughout human pregnancy | Homo sapiens |
| SDY1290 | Dendritic Cell Maturation Dynamics | Homo sapiens |
| SDY1302 | Prematurity, Respiratory outcomes, Immune System, and Microbiome Study (PRISM) | Homo sapiens |
| SDY1256 | Small Sample- Big Data: The Gambia | Homo sapiens |

| | | |
|---|---|---|
| SDY1288 | Chikungunya virus infection immunoprofiling in pediatric cohort | Homo sapiens |
| SDY903 | Human Immune Signature of Zika virus infection | Homo sapiens |
| SDY1369 | Immune signatures of responses to infection with Dengue and Zika virus | Homo sapiens |
| SDY1371 | Profiling of NK cell ligands on monocytes infected with Influenza A virus | Homo sapiens |
| SDY1634 | Charge-Altering Releasable Transporters Enable Specific Phenotypic Manipulation Of Resting Primary Natural Killer Cells | Homo sapiens |
| SDY1385 | IFN-g-independent immune markers of Mycobacterium tuberculosis exposure | Homo sapiens |
| SDY1335 | Immune Profiling Assay Development for Pertussis vaccines | Homo sapiens |
| SDY1389 | Lymph node reservoirs for long-lived memory T cells | Homo sapiens |
| SDY1086 | Responses to Inactivated Influenza Vaccine (IIV) in adults with or without antibiotics | Homo sapiens |
| SDY1535 | TIGIT is upregulated by HIV-1 infection and marks a highly functional adaptive and mature subset of natural killer cells | Homo sapiens |
| SDY1555 | Allele-specific expression changes dynamically during T cell activation in HLA and other autoimmune loci | Homo sapiens |
| SDY1597 | Immune Cell Repertoires in Breast Cancer Patients after Adjuvant Chemotherapy | Homo sapiens |
| SDY1603 | Investigating the natural killer cell response to acute dengue infection. | Homo sapiens |
| SDY1620 | Treated HIV infection induces alterations in phenotype but not HIV-specific function of peripheral blood natural killer cells | Homo sapiens |
| SDY1625 | Double Expressor Assessment | Homo sapiens |
| SDY1630 | Effects of tissue localization on Natural Killer (NK) cell phenotypic and functional diversity | Homo sapiens |
| SDY1658 | A comparative CyTOF analysis of resected-tumor samples from human GBM patients and mouse GBM-tumors (see companion study SDY1637) | Homo sapiens |
| SDY1655 | Longitudinal Analyses Reveal Immunological Misfiring in Severe COVID-19 (Companion study to SDY1648) | Homo sapiens |
| SDY1886 | Multiscale PHATE identifies multimodal signatures of COVID-19 | Homo sapiens |
| SDY1666 | KSHV tropism in B lymphocytes | Homo sapiens |
| SDY1667 | Cross-reactive SARS-CoV-2 T cell epitopes in unexposed humans | Homo sapiens |

| | | |
|---|---|---|
| SDY1640 | T and B cell responses to SARS-CoV-2 coronavirus | Homo sapiens |
| SDY1680 | Comorbid illnesses are associated with altered adaptive immune responses to SARS-CoV-2 | Homo sapiens |
| SDY1647 | CD16 is upregulated in HIV-exposed seronegative women and may mediate enhanced antibody-dependent cytotoxicity | Homo sapiens |
| SDY1734 | Controlled malaria infection in Europeans and Africans | Homo sapiens |
| SDY1600 | Mapping Immune Responses to CMV in Renal Transplantation | Homo sapiens |
| SDY1708 | Broad dysregulation of innate immunity and hematopoiesis distinguishes mild from severe COVID-19 | Homo sapiens |
| SDY1743 | Antigen-Specific Adaptive Immunity to SARS-CoV-2 in Acute COVID-19 | Homo sapiens |
| SDY1773 | Human plasmacytoid dendritic cells mount a distinct antiviral response to virus-infected cells | Homo sapiens |
| SDY1767 | Longitudinal profiling of respiratory and systemic immune responses in severe COVID-19 | Homo sapiens |
| SDY1844 | Natural killer cell repertoire and function in HIV-1 persistence | Homo sapiens |
| SDY1870 | Kinetics of immunological memory to SARS-CoV-2 | Homo sapiens |
| SDY1885 | Heterogeneity of human anti-viral immunity shaped by virus, tissue, age, and sex | Homo sapiens |
| SDY1944 | Flow cytometer data for investigation into an integrated microfluidic system for basophil isolation from whole blood | Homo sapiens |
| SDY1669 | Mild and severe COVID-19 | Homo sapiens |
| SDY1961 | Age-related signs of immunosenescence correlate with 3 poor outcome of mRNA COVID-19 vaccination in older 4 adults | Homo sapiens |
| SDY165 | Characterization of in vitro Stimulated B Cells from Human Subjects | Homo sapiens |
| SDY217 | Orchestration of CD4 T cell epitope preferences after multi-peptide immunization | Mus musculus |
| SDY579 | Assessing the role of NLRX1 during mucosal immune responses to H. pylori in mice | Mus musculus |
| SDY598 | HP45 | Mus musculus |
| SDY601 | HP47 | Mus musculus |
| SDY583 | Immune Responses to Seasonal Influenza Vaccination in Mouse model | Mus musculus |
| SDY585 | 2013 Murine Vaccine with Adjuvant Study | Mus musculus |
| SDY827 | An Interactive Reference Framework for Modeling a Dynamic Immune System | Mus musculus |

| | | |
|---|---|---|
| SDY901 | Activation of the PD-1 Pathway Contributes to Immune Escape in EGFR-Driven Lung Tumors | Mus musculus |
| SDY1108 | Systemic Immunity for Cancer Immunotherapy | Mus musculus |
| SDY1176 | Extensive homeostatic T cell phenotypic variation within the Collaborative Cross | Mus musculus |
| SDY1502 | Platelets attach to lung ILC2 expressing PSGL-1 and influence ILC2 function | Mus musculus |
| SDY1595 | MDR1 expression in hematopeotic cells | Mus musculus |
| SDY1536 | Unconventional ST2- and CD127-negative lung ILC2 populations are induced by the fungal allergen Alternaria | Mus musculus |
| SDY1618 | Superior mouse eosinophil depletion in vivo targeting transgenic Siglec-8 instead of endogenous Siglec-F: mechanisms and pitfalls | Mus musculus |
| SDY1631 | The roles of CD103 and CD49a in adherence and motility of CD8 T cells | Mus musculus |
| SDY1637 | Immune phenotyping of diverse syngeneic murine brain tumors identifies immunologically distinct types (see companion study SDY1658) | Mus musculus |
| SDY1774 | Tumor Immune Microenvironment of mIDH1 and wtIDH1 Glioma GEMMs | Mus musculus |
| SDY139 | The peptide specificity of the endogenous T follicular helper cell repertoire generated after protein immunization | Mus musculus |
| SDY1424 | MaHPIC: Host Aotus nancymaae infected with P. vivax Brazil VII | Aotus nancymaae |
| SDY1015 | MaHPIC: Host M. mulatta infected with P. cynomolgi | Macaca mulatta |
| SDY1409 | MaHPIC: Host M. mulatta infected with homologous and heterologous strains of P. cynomolgi | Macaca mulatta |
| SDY1411 | MaHPIC: Host M. mulatta infected with P. coatneyi | Macaca mulatta |

Table S11. Summary of the inputted and matched marker names from the bone marrow study published by Samusik et al[7]. 'Inputted' refers to the marker name as described by the study, while 'matched' refers to the marker name after standardization and automatic matching within the Part 2 workflow. The matched PRO or GO terms were used for cell type matching in the CL. Only PRO or GO terms that were included in the CL at the time of analysis are included.

| **Inputted Marker Name** | **Matched Marker Name** | **Matched PRO/GO Term** |
|---|---|---|
| B220 | B220 | http://purl.obolibrary.org/obo/PR_000001014 |
| CD115 | CD115 | http://purl.obolibrary.org/obo/PR_000002062 |
| CD11B | CD11B | http://purl.obolibrary.org/obo/PR_000001012 |
| CD11C | CD11C | http://purl.obolibrary.org/obo/PR_000001013 |
| CD138 | CD138 | http://purl.obolibrary.org/obo/PR_000001935 |
| CD16_32 | CD16<br>CD32 | http://purl.obolibrary.org/obo/PR_000001483<br>http://purl.obolibrary.org/obo/PR_000001479<br>http://purl.obolibrary.org/obo/PR_000001481 |
| CD19 | CD19 | http://purl.obolibrary.org/obo/PR_000001002 |
| CD34 | CD34 | http://purl.obolibrary.org/obo/PR_000001003 |
| CD3 | CD3E | http://purl.obolibrary.org/obo/PR_000001020 |
| CD4 | CD4 | http://purl.obolibrary.org/obo/PR_000001004 |
| CD43 | CD43 | http://purl.obolibrary.org/obo/PR_000001879 |
| CD49B | CD49B | http://purl.obolibrary.org/obo/PR_000001008 |
| CD64 | CD64 | http://purl.obolibrary.org/obo/PR_000001465 |
| CD8 | CD8A<br>CD8ALPHABETA | http://purl.obolibrary.org/obo/PR_000001084<br>http://purl.obolibrary.org/obo/PR_000025402 |
| CKIT | CKIT | http://purl.obolibrary.org/obo/PR_000002065 |
| F480 | EMR1 | http://purl.obolibrary.org/obo/PR_000001813 |
| FCER1A | FCER1A | http://purl.obolibrary.org/obo/PR_000007431 |
| IGD | IGD | http://purl.obolibrary.org/obo/GO_0071738 |
| IGM | IGM | http://purl.obolibrary.org/obo/GO_0071753 |
| LY6C | LY6C | http://purl.obolibrary.org/obo/PR_000002980 |
| LY6G | LY6G | http://purl.obolibrary.org/obo/PR_000002978 |
| MHCII | MHCII | http://purl.obolibrary.org/obo/GO_0042613 |
| NKP46 | NKP46 | http://purl.obolibrary.org/obo/PR_000001893 |
| SCA1 | SCA1 | http://purl.obolibrary.org/obo/PR_000002979 |
| SIGLECF | SIGLECF | http://purl.obolibrary.org/obo/PR_000001927 |
| TCRB | TCRAB | http://purl.obolibrary.org/obo/GO_0042105 |
| TCRGD | TCRGD | http://purl.obolibrary.org/obo/GO_0042106 |
| 120G8 | Not matched | |

Table S12. Summary of the inputted and matched marker names from the PBMC study published by Kimmey et al[8]. ‘Inputted’ refers to the marker name as described by the study, while ‘matched’ refers to the marker name after standardization and automatic matching within the Part 2 workflow. The matched PRO or GO terms were used for cell type matching in the CL. Only PRO or GO terms that were included in the CL at the time of analysis are included.

| **Inputted Marker Name** | **Matched Marker Name** | **Matched PRO/GO Term** |
|---|---|---|
| CD11C | CD11C | http://purl.obolibrary.org/obo/PR_000001013 |
| CD123 | CD123 | http://purl.obolibrary.org/obo/PR_000001865 |
| CD14 | CD14 | http://purl.obolibrary.org/obo/PR_000001889 |
| CD19 | CD19 | http://purl.obolibrary.org/obo/PR_000001002 |
| CD20 | CD20 | http://purl.obolibrary.org/obo/PR_000001289 |
| CD3 | TCR<br>TCRAB<br>TCRGD<br>CD3E | http://purl.obolibrary.org/obo/GO_0042101<br>http://purl.obolibrary.org/obo/GO_0042105<br>http://purl.obolibrary.org/obo/GO_0042106<br>http://purl.obolibrary.org/obo/PR_000001020 |
| CD38 | CD38 | http://purl.obolibrary.org/obo/PR_000001408 |
| CD4 | CD4 | http://purl.obolibrary.org/obo/PR_000001004 |
| CD56 | CD56 | http://purl.obolibrary.org/obo/PR_000001024 |
| HLADR | HLADR | http://purl.obolibrary.org/obo/GO_0042613 |
| CD16 | Not included due to ‘null’ assignment in Part 1 | |

Table S13. Summary of the inputted and matched marker names from the unpublished spectral cytometry PBMC study. 'Inputted' refers to the marker name as described by the study, while 'matched' refers to the marker name after standardization and automatic matching within the Part 2 workflow. The matched PRO or GO terms were used for cell type matching in the CL. Only PRO or GO terms that were included in the CL at the time of analysis are included.

| **Inputted Marker Name** | **Matched Marker Name** | **Matched PRO/GO Term** |
|---|---|---|
| CCR7 | CCR7 | http://purl.obolibrary.org/obo/PR_000001203 |
| CD19 | CD19 | http://purl.obolibrary.org/obo/PR_000001002 |
| CD3 | TCR<br>TCRAB<br>TCRGD<br>CD3E | http://purl.obolibrary.org/obo/GO_0042101<br>http://purl.obolibrary.org/obo/GO_0042105<br>http://purl.obolibrary.org/obo/GO_0042106<br>http://purl.obolibrary.org/obo/PR_000001020 |
| CD4 | CD4 | http://purl.obolibrary.org/obo/PR_000001004 |
| CD45RA | CD45RA | http://purl.obolibrary.org/obo/PR_000001015 |
| CD56 | CD56 | http://purl.obolibrary.org/obo/PR_000001024 |
| CD8 | CD8A<br>CD8ALPHABETA | http://purl.obolibrary.org/obo/PR_000001084<br>http://purl.obolibrary.org/obo/PR_000025402 |
| CD14 | Not included due to 'null' assignment in Part 1 | |

Table S14. Summary of the inputted and matched marker names from OMIP 54[9]. ‘Inputted’ refers to the marker name as described by the study, while ‘matched’ refers to the marker name after standardization and automatic matching within the Part 2 workflow. The matched PRO or GO terms were used for cell type matching in the CL. Only PRO or GO terms that were included in the CL at the time of analysis are included.

| **Inputted Marker Name** | **Matched Marker Name** | **Matched PRO/GO Term** |
|---|---|---|
| CCR2 | CCR2 | http://purl.obolibrary.org/obo/PR_000001199 |
| CD11B | CD11B | http://purl.obolibrary.org/obo/PR_000001012 |
| CD11C | CD11C | http://purl.obolibrary.org/obo/PR_000001013 |
| CD3 | TCR<br>TCRAB<br>TCRGD<br>CD3E | http://purl.obolibrary.org/obo/GO_0042101<br>http://purl.obolibrary.org/obo/GO_0042105<br>http://purl.obolibrary.org/obo/GO_0042106<br>http://purl.obolibrary.org/obo/PR_000001020 |
| CD4 | CD4 | http://purl.obolibrary.org/obo/PR_000001004 |
| CD44 | CD44 | http://purl.obolibrary.org/obo/PR_000001307 |
| CD49D | CD49D | http://purl.obolibrary.org/obo/PR_000009129 |
| CD8 | CD8A<br>CD8ALPHABETA | http://purl.obolibrary.org/obo/PR_000001084<br>http://purl.obolibrary.org/obo/PR_000025402 |
| FOXP3 | FOXP3 | http://purl.obolibrary.org/obo/PR_000001350 |
| LY6C | LY6C | http://purl.obolibrary.org/obo/PR_000002980 |
| LY6G | LY6G | http://purl.obolibrary.org/obo/PR_000002978 |
| TCRB | TCRAB | http://purl.obolibrary.org/obo/GO_0042105 |

Table S15. Summary of the inputted and matched marker names from OMIP 63[10]. 'Inputted' refers to the marker name as described by the study, while 'matched' refers to the marker name after standardization and automatic matching within the Part 2 workflow. The matched PRO or GO terms were used for cell type matching in the CL. Only PRO or GO terms that were included in the CL at the time of analysis are included. Note that there was no exact equivalent to TCR Vα24JαQ or TCR Vα7.2 within PRO, so the more general TCR marker was used instead.

| Inputted Marker Name | Matched Marker Name | Matched PRO/GO Term |
|---|---|---|
| CCR6 | CCR6 | http://purl.obolibrary.org/obo/PR_000001202 |
| CCR7 | CCR7 | http://purl.obolibrary.org/obo/PR_000001203 |
| CD10 | CD10 | http://purl.obolibrary.org/obo/PR_000001898 |
| CD127 | CD127 | http://purl.obolibrary.org/obo/PR_000001869 |
| CD161 | CD161 | http://purl.obolibrary.org/obo/PR_000025670 |
| CD20 | CD20 | http://purl.obolibrary.org/obo/PR_000001289 |
| CD25 | CD25 | http://purl.obolibrary.org/obo/PR_000001380 |
| CD27 | CD27 | http://purl.obolibrary.org/obo/PR_000001963 |
| CD3E | CD3E | http://purl.obolibrary.org/obo/PR_000001020 |
| CD4 | CD4 | http://purl.obolibrary.org/obo/PR_000001004 |
| CD45RA | CD45RA | http://purl.obolibrary.org/obo/PR_000001015 |
| CD56 | CD56 | http://purl.obolibrary.org/obo/PR_000001024 |
| CD8 | CD8A<br>CD8ALPHABETA | http://purl.obolibrary.org/obo/PR_000001084<br>http://purl.obolibrary.org/obo/PR_000025402 |
| CXCR3 | CXCR3 | http://purl.obolibrary.org/obo/PR_000001207 |
| CXCR5 | CXCR5 | http://purl.obolibrary.org/obo/PR_000001209 |
| IGA | IGA | http://purl.obolibrary.org/obo/GO_0071745 |
| IGG | IGG | http://purl.obolibrary.org/obo/GO_0071735 |
| IGM | IGM | http://purl.obolibrary.org/obo/GO_0071753 |
| TCR VAB | TCRAB | http://purl.obolibrary.org/obo/GO_0042105 |
| TCR VGD | TCRGD | http://purl.obolibrary.org/obo/GO_0042106 |
| TCR Vα24JαQ | TCR | http://purl.obolibrary.org/obo/GO_0042101 |
| TCR Vα7.2 | TCR | http://purl.obolibrary.org/obo/GO_0042101 |
| CD161 | Matched to PRO term, but not in the CL | |

Table S16. Summary of the inputted and matched marker names from OMIP 78[11]. 'Inputted' refers to the marker name as described by the study, while 'matched' refers to the marker name after standardization and automatic matching within the Part 2 workflow. The matched PRO or GO terms were used for cell type matching in the CL. Only PRO or GO terms that were included in the CL at the time of analysis are included.

| **Inputted Marker Name** | **Matched Marker Name** | **Matched PRO/GO Term** |
|---|---|---|
| CD11B | CD11B | http://purl.obolibrary.org/obo/PR_000001012 |
| CD11C | CD11C | http://purl.obolibrary.org/obo/PR_000001013 |
| CD123 | CD123 | http://purl.obolibrary.org/obo/PR_000001865 |
| CD133 | CD133 | http://purl.obolibrary.org/obo/PR_000001786 |
| CD138 | CD138 | http://purl.obolibrary.org/obo/PR_000001935 |
| CD14 | CD14 | http://purl.obolibrary.org/obo/PR_000001889 |
| CD141 | CD141 | http://purl.obolibrary.org/obo/PR_000002105 |
| CD16 | CD16 | http://purl.obolibrary.org/obo/PR_000001483 |
| CD19 | CD19 | http://purl.obolibrary.org/obo/PR_000001002 |
| CD1C | CD1C | http://purl.obolibrary.org/obo/PR_000002027 |
| CD24 | CD24 | http://purl.obolibrary.org/obo/PR_000001932 |
| CD27 | CD27 | http://purl.obolibrary.org/obo/PR_000001963 |
| CD3 | TCR<br>TCRAB<br>TCRGD<br>CD3E | http://purl.obolibrary.org/obo/GO_0042101<br>http://purl.obolibrary.org/obo/GO_0042105<br>http://purl.obolibrary.org/obo/GO_0042106<br>http://purl.obolibrary.org/obo/PR_000001020 |
| CD303 | CD303 | http://purl.obolibrary.org/obo/PR_000001292 |
| CD34 | CD34 | http://purl.obolibrary.org/obo/PR_000001003<br>http://purl.obolibrary.org/obo/PR_P28906 |
| CD38 | CD38 | http://purl.obolibrary.org/obo/PR_000001408 |
| CD4 | CD4 | http://purl.obolibrary.org/obo/PR_000001004 |
| CD45RO | CD45RO | http://purl.obolibrary.org/obo/PR_000001017 |
| CD56 | CD56 | http://purl.obolibrary.org/obo/PR_000001024 |
| CD8 | CD8A<br>CD8ALPHABETA | http://purl.obolibrary.org/obo/PR_000001084<br>http://purl.obolibrary.org/obo/PR_000025402 |
| HLADR | HLADR | http://purl.obolibrary.org/obo/GO_0042613 |
| IGD | IGD | http://purl.obolibrary.org/obo/GO_0071738 |
| IGG | IGG | http://purl.obolibrary.org/obo/GO_0071735 |
| IGM | IGM | http://purl.obolibrary.org/obo/GO_0071753 |
